# Advanced Whole Genome Sequencing Using an Entirely PCR-free Massively Parallel Sequencing Workflow

**DOI:** 10.1101/2019.12.20.885517

**Authors:** Hanjie Shen, Pengjuan Liu, Zhanqing Li, Fang Chen, Hui Jiang, Shiming Shi, Yang Xi, Qiaoling Li, Xiaojue Wang, Jing Zhao, Xinming Liang, Yinlong Xie, Lin Wang, Wenlan Tian, Tam Berntsen, Andrei Alexeev, Yinling Luo, Meihua Gong, Jiguang Li, Chongjun Xu, Nina Barua, Snezana Drmanac, Sijie Dai, Zilan Mi, Han Ren, Zhe Lin, Ao Chen, Wenwei Zhang, Feng Mu, Xun Xu, Xia Zhao, Yuan Jiang, Radoje Drmanac

## Abstract

**Background:** Systematic errors can be introduced from DNA amplification during massively parallel sequencing (MPS) library preparation and sequencing array formation. Polymerase chain reaction (PCR)-free genomic library preparation methods were previously shown to improve whole genome sequencing (WGS) quality on the Illumina platform, especially in calling insertions and deletions (InDels). We hypothesized that substantial InDel errors continue to be introduced by the remaining PCR step of DNA cluster generation. In addition to library preparation and sequencing, data analysis methods are also important for the accuracy of the output data.In recent years, several machine learning variant calling pipelines have emerged, which can correct the systematic errors from MPS and improve the data performance of variant calling.

**Results:** Here, PCR-free libraries were sequenced on the PCR-free DNBSEQ™ arrays from MGI Tech Co., Ltd. (referred to as MGI) to accomplish the first true PCR-free WGS which the whole process is truly not only PCR-free during library preparation but also PCR-free during sequencing. We demonstrated that PCR-based WGS libraries have significantly (about 5 times) more InDel errors than PCR-free libraries.Furthermore, PCR-free WGS libraries sequenced on the PCR-free DNBSEQ™ platform have up to 55% less InDel errors compared to the NovaSeq platform, confirming that DNA clusters contain PCR-generated errors.In addition, low coverage bias and less than 1% read duplication rate was reproducibly obtained in DNBSEQ™ PCR-free using either ultrasonic or enzymatic DNA fragmentation MGI kits combined with MGISEQ-2000. Meanwhile, variant calling performance (single-nucleotide polymorphisms (SNPs) F-score>99.94%, InDels F-score>99.6%) exceeded widely accepted standards using machine learning (ML) methods (DeepVariant or DNAscope).

**Conclusions:** Enabled by the new PCR-free library preparation kits, ultra high-thoughput PCR-free sequencers and ML-based variant calling, true PCR-free DNBSEQ™ WGS provides a powerful solution for improving WGS accuracy while reducing cost and analysis time, thus facilitating future precision medicine, cohort studies, and large population genome projects.

## Background

Massively parallel sequencing (MPS, also known as next-generation sequencing (NGS)) technology has revolutionized basic biology and precision medicine during the past decade. There is an increasing clinical demand for whole genome sequencing (WGS) to be used as a single test, especially in conditions when partially sequencing the genome via targeted panels such as whole-exome sequencing (WES) or target region sequencing could potentially fail to detect all pathogenic variants in a large fraction of Mendelian disorder cases [1,2,3]. A variety of studies have already demonstrated the feasibility of WGS for investigating rare diseases [4], cancers [5], and infectious diseases [6]. More importantly,WGS currently costs no more than one thousand dollars[7], has faster turnaround time, and accomplishes greater depth, making it more economical to conduct large-scale projects.

The standard WGS workflow includes MPS library preparation, clonal DNA array (e.g., PCR clusters) generation, on-chip sequencing, read filtering, mapping, and variant calling (also known as secondary analysis). Many efforts have been undertaken to further reduce the cost and turnaround time while improving WGS data performance. For example, an optimized WGS library protocol that eliminates the need for PCR—PCR-free WGS—has been developed to eliminate the amplification bias (including coverage bias, GC bias [8,9,10,11,12]), copy errors (mainly refers to InDels errors [13]) and duplication reads [14] during library preparation process. Another benefit of excluding the PCR step from WGS library preparation is significantly shorter turnaround time and lower cost.

In addition to the new library construction chemistry, different innovative bioinformatics tools have been applied to expedite data analysis without sacrificing accuracy. The steps of a standard analysis workflow are typically as follows: 1) trim and filter read data; 2) align raw data to a reference genome, 3) call germline or somatic variants, and 4) conduct tertiary analysis and generate reports. The pipeline for germline short variant discovery developed at the Broad Institute with the Genome Analysis Toolkit (GATK) [15] is currently the industry standard for variant calling on WGS. However, the traditional GATK pipeline takes approximately two days for whole genome data processing on a standard 24-thread machine [16]. To achieve better detection accuracy and account for systematic errors in the WGS workflow, researchers have explored machine learning and deep learning-based algorithms and developed several new analysis pipelines such as GATK CNNScoreVariants from the Broad Institute [17], Deepvariant from the Google Brain team [18], DNAscope from Sentieon [19], and Clairvoyante from R. Luo et al. [20].

In this report, we present an entirely PCR-free MPS workflow by constructing PCR-free WGS libraries and sequencing them on MGI’s PCR-free DNBSEQ™ arrays. Both PCR-free WGS sets (mechanical fragmentation with MGIEasy PCR-Free DNA Library Prep Set or enzymatic fragmentation with MGIEasy FS PCR-Free DNA Library Prep Set) demonstrated highly reproducible data quality with as little as 200 ng or 50 ng genomic DNA as input. More importantly, a significant improvement was achieved in InDel calling with GATK from an average F-score of 95.43% in three PCR libraries to 99.05% in three PCR-free WGS libraries. By incorporating machine learning-based algorithms, the F-score of 15x PCR-free libraries analyzed with DNAscope or DeepVariant can outperform that of 30x PCR-prepared libraries analyzed with GATK in some scenarios. As hypothesized, complete PCR-free WGS significantly decreased false positive (FP) and false negative (FN) InDel calls, leading to better InDel calling accuracy (99.3-99.6% F-score) and up to 55% less InDel errors than WGS of Illumina’s PCR-free libraries sequenced on a Novaseq array of PCR-generated DNA clusters (F-score 98.0-99.3%), even with less genomic DNA input. Additionally, a very low duplication rate of less than 1% was achieved in the DNBSEQ™ PCR-free WGS, resulting in more informative reads and further reducing sequencing cost. In summary, the advanced PCR-free WGS reported herein could lead to wider adoption of WGS in genomic research and gradually be incorporated into clinical practice for the urgent diagnosis of rare disease and for improved disease prevention.

## Results

### An entirely PCR-free MPS workflow

PCR is frequently used to increase template quantity during MPS library construction. PCR is also an essential step of “bridge amplification” to generate identical copies on a flow cell surface [21]. Here, we describe an MPS workflow called DNBSEQ™ PCR-free, which completely eliminates PCR amplification during both library and array preparation (Fig. 1a). DNBSEQ™ PCR-free starts with DNA fragmentation, which is followed by size selection using solid-phase reversible immobilization (SPRI) beads. A single-tube protocol is used to conduct multiple sequential enzymatic reactions and attach a barcoded adapter to the DNA of interest. After removing excess adapters, ssCir DNA are formed, and these serve as template in rolling circle replication (RCR) for DNB preparation. DNBs are then loaded onto patterned flow slides and sequenced [22]. In contrast to bridge amplification, RCR is a linear amplification from the original ssCir template, and therefore clonal errors cannot be generated [23,24], unlike bridge PCR- or emulsion PCR-based preparation of sequencing arrays. PCR-free DNBSEQ™ completely avoids PCR errors in template amplification and library cloning and faithfully preserves the original landscape of the genome with rare index mis-assignment [25].

**Figure 1.**
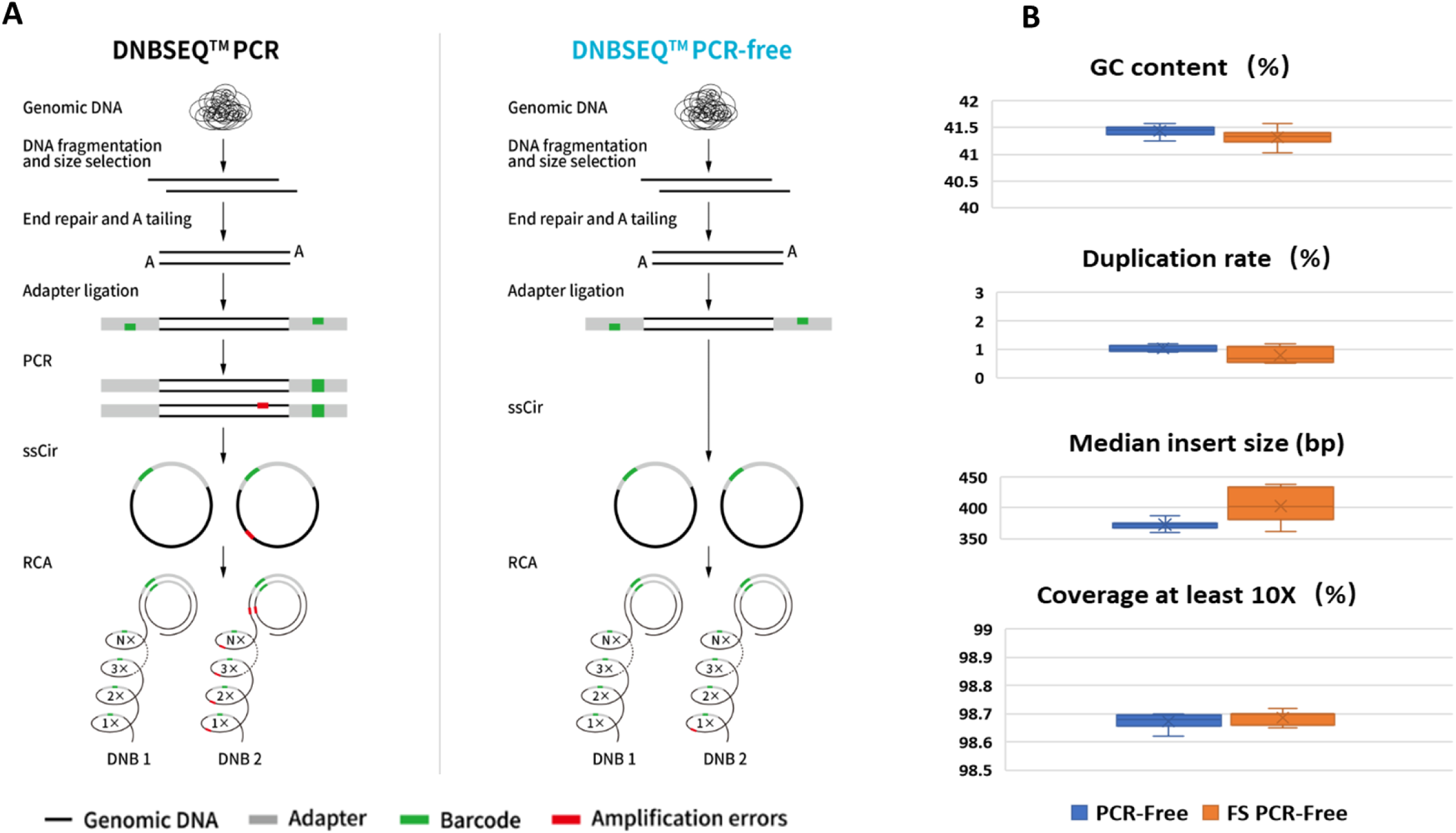
DNBSEQ™ WGS PCR vs. PCR-free workflows and general performance of PCR-free libraries. (a) MPS library construction workflows of WGS PCR and PCR-free libraries. Rolling circle amplification (RCA) is used to increase signal intensity during array formation, which is followed by sequencing DNBs with DNBSEQ™ technology. Individual copies from the same DNB are replicated independently using the same ssCir template. Therefore, amplification errors cannot accumulate. Black, genomic DNA; gray rectangle, adapter; green, barcode; red, amplification errors. (b) Two sets of 9 replicates from 1 μg NA12878 DNA were processed with MGIEasy PCR-Free DNA Library Prep Set (blue) or MGIEasy FS PCR-Free DNA Library Prep Set (orange). The GC content, Duplication rate, Median Insert size, and regions with >10x Coverage were calculated and plotted. The error bars represent the standard deviations across the replicates.

Two sets (MGIEasy PCR-Free DNA Library Pre Set and MGIEasy FS PCR-Free DNA Library Pre Set) from MGI are used to prepare the DNBSEQ™ PCR-free libraries. The PCR-Free set is used with ultrasonic fragmented samples, whereas the FS PCR-Free set includes sequential enzymatic fragmentation reactions, end repair/dA-tailing, and adapter addition in a single tube. We compared the performance of both sets with nine libraries constructed from 1 μg NA12878 reference genomic DNA and sequenced with paired-end 150-bp read length. Fig.1b summarizes the QC statistics, including GC content, duplication rate, median insert size, and regions with > 10x coverage. Both sets showed highly reproducible performance; the PCR-Free set showed slightly less variation compared with the FS PCR-Free set. We also tested different input quantities (1 μg, 500 ng, and 200 ng for both sets and 50 ng only for the FS PCR-Free Set) and observed comparable performance (Supplementary Fig. 1S).

### Minimal GC bias for genomes with different GC content

The relationship between GC content and read coverage across a genome, known as GC bias, can be greatly affected by MPS library preparation, cluster/array amplification, and sequencing. To evaluate the performance of the DNBSEQ™ PCR-free MPS workflow with regard to GC bias, DNA samples from bacterial strains with GC contents of 38% and 62% were processed with the PCR-Free and FS PCR-Free sets mentioned above. Libraries were prepared according to kit instructions and sequenced on MGISEQ-2000 with paired-end sequencing (2×150 bp). Fig.2 shows the number of reads covering different regions normalized by the mean read coverage and plotted against the genomic GC content of the corresponding region. Normalized coverage lower or higher than 1 indicates certain GC bias. The high-GC and low-GC genomes demonstrated fairly even coverage of reads across the genome with either ultrasonic shearing or enzymatic shearing. Overall, the DNBSEQ™ PCR-free MPS workflow demonstrated minimal GC bias in genomes with significantly varied GC content.

**Figure 2.**
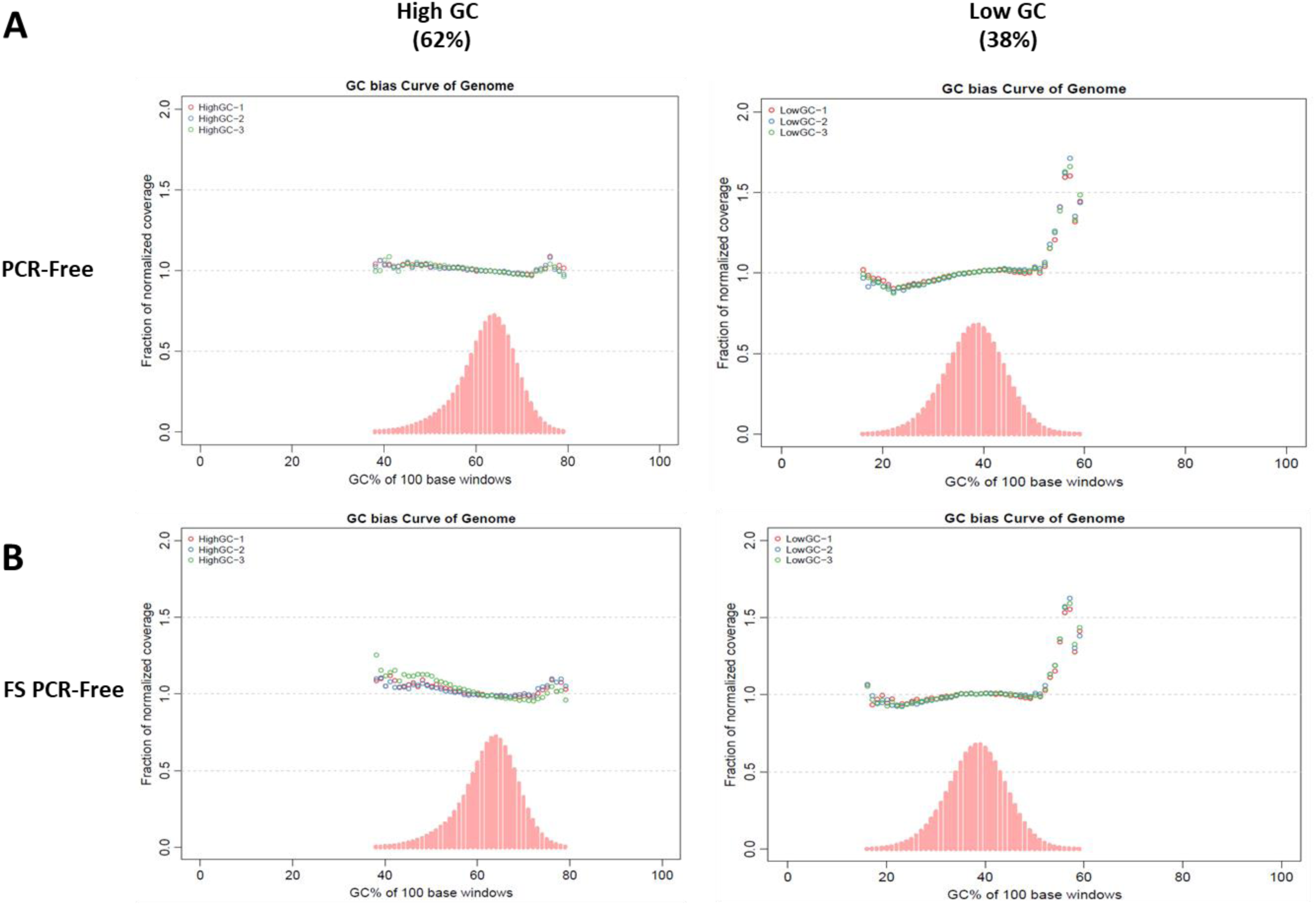
Coverage of the microbial genomes, *Olsenella profusa* (62% GC, left) and *Bacillus megaterium* (38% GC, right) with (A) MGIEasy PCR-Free DNA Library Pre Set and (B) MGIEasy FS PCR-Free DNA Library Pre Set. Read coverage across the range of the GC content, calculated in 100-bp windows (pink bars) and normalized coverage (colored dots). Three replicates (red, blue, and green dots) were included in the normalized coverage vs. GC content analysis.

### Low duplicate rate and high variant calling F-score for GIAB with DNBSEQ™ PCR-free

We compared the sequencing accuracy of MPS libraries prepared with or without PCR using DNBSEQ™ sequencing technology [26]. After ultrasonic fragmentation, NA12878 DNA was processed according to the instructions of the MGIEasy PCR-Free DNA Library Prep Set to construct three PCR-free WGS libraries. Three PCR-based WGS libraries were also prepared from the same DNA sample. Each library was sequenced individually on one lane of MGISEQ-2000 with paired-end sequencing (2×150 bp) (see Methods). The total raw data of each lane was greater than 120G with GC content ranging from 41.46% to 41.78% (Table 1).

**Table 1.** Mapping performance of three replicates for PCR and PCR-free libraries. QC matrix from 3 PCR and 3 PCR-free libraries was collected. Each sample was down-sampled to 15x and 30x in addition to full lane (FL, approximately 46x coverage).

| Method | PCR-1 |  |  | PCR-2 |  |  | PCR-3 |  |  | PCR-free-1 |  |  | PCR-free-2 |  |  | PCR-free-3 |  |  |
| --- | --- | --- | --- | --- | --- | --- | --- | --- | --- | --- | --- | --- | --- | --- | --- | --- | --- | --- |
| Depth | 15x | 30x | FL | 15x | 30x | FL | 15x | 30x | FL | 15x | 30x | FL | 15x | 30x | FL | 15x | 30x | FL |
| Clean reads (x10<sup>6</sup>) | 296.64 | 633.73 | 940.37 | 296.80 | 633.76 | 954.92 | 296.45 | 632.78 | 949.34 | 295.93 | 631.36 | 926.38 | 296.03 | 632.98 | 944.92 | 296.13 | 631.49 | 964.32 |
| Clean bases (Gb) | 44.49 | 95.05 | 141.05 | 44.52 | 95.06 | 143.23 | 44.46 | 94.91 | 142.40 | 44.39 | 94.70 | 138.95 | 44.40 | 94.94 | 141.73 | 44.42 | 94.72 | 144.64 |
| Insert size peak (bp) | 383 | 383 | 383 | 374 | 374 | 374 | 375 | 375 | 375 | 374 | 374 | 375 | 375 | 375 | 375 | 375 | 375 | 375 |
| GC content (%) | 41.64 | 41.64 | 41.64 | 41.78 | 41.78 | 41.78 | 41.49 | 41.49 | 41.49 | 41.53 | 41.53 | 41.53 | 41.48 | 41.48 | 41.48 | 41.46 | 41.47 | 41.46 |
| Mapping rate (%) | 99.91 | 99.82 | 99.82 | 99.91 | 99.84 | 99.84 | 99.89 | 99.82 | 99.82 | 99.89 | 99.83 | 99.83 | 99.72 | 99.65 | 99.65 | 99.93 | 99.86 | 99.86 |
| Unique rate (%) | 99.42 | 95.41 | 94.78 | 99.48 | 95.53 | 94.92 | 99.35 | 95.23 | 94.51 | 99.69 | 95.76 | 95.38 | 99.71 | 95.82 | 95.44 | 99.77 | 95.95 | 95.61 |
| Duplicate rate (%) | 0.58 | 1.56 | 2.24 | 0.52 | 1.44 | 2.10 | 0.65 | 1.72 | 2.49 | 0.31 | 1.07 | 1.50 | 0.29 | 1.03 | 1.45 | 0.23 | 0.91 | 1.31 |
| Mismatch rate (%) | 0.89 | 0.87 | 0.86 | 0.91 | 0.89 | 0.88 | 0.95 | 0.93 | 0.92 | 0.74 | 0.74 | 0.73 | 0.74 | 0.74 | 0.73 | 0.71 | 0.70 | 0.70 |
| Average seq depth (X) | 15.37 | 30.83 | 45.46 | 15.39 | 30.89 | 46.27 | 15.34 | 30.71 | 45.74 | 15.37 | 30.81 | 45.04 | 15.35 | 30.86 | 45.91 | 15.4 | 30.91 | 47.04 |
| Coverage (%) | 99.07 | 99.13 | 99.16 | 99.07 | 99.13 | 99.16 | 99.08 | 99.14 | 99.17 | 99.1 | 99.16 | 99.18 | 99.1 | 99.16 | 99.18 | 99.1 | 99.16 | 99.18 |
| Coverage at least 4X (%) | 98.65 | 98.96 | 99.03 | 98.64 | 98.97 | 99.03 | 98.65 | 98.97 | 99.04 | 98.76 | 99.01 | 99.06 | 98.75 | 99.01 | 99.06 | 98.74 | 99.00 | 99.06 |
| Coverage at least 10X (%) | 89.78 | 98.58 | 98.83 | 89.61 | 98.58 | 98.84 | 89.55 | 98.58 | 98.84 | 90.5 | 98.70 | 98.89 | 90.44 | 98.69 | 98.89 | 90.49 | 98.69 | 98.90 |

The raw reads were down-sampled from the original full lane (approximately 46x) to create additional 30x and 15x depth datasets. After raw read filtering, clean reads were aligned to the human reference genome with decoy sequence hs37d5. The mapping quality matrix is summarized in Table 1. We compared all 3 dataset depths from the 6 libraries (18 datasets in total). In 30x depth data, WGS PCR and PCR-free libraries had a similarly high mapping rate of 99.8% and overall coverage greater than 99.1%. PCR-free libraries showed slightly lower duplication and mismatch rates of approximately 1% and 0.7%, respectively on average, whereas these values were 1.5% and 0.9% in PCR libraries. Other depth datasets showed similar patterns. Theoretically, because of the true PCR-free workflow, the duplicate rate should be zero, but it is possible that the same DNB is read twice or more due to optical overflow (optical duplicates) or that the same regions from different genome copies produced in the DNA fragmentation step (natural duplicates) are sequenced and incorrectly marked as duplicates by QC tools [27,28].

We next used the three variant callers GATK, DeepVariant, and DNAscope (see Methods) to assess the accuracy of PCR and PCR-free methods (Table 2, Supplementary Table 1S, and Fig. 3). The GATK HaplotypeCaller has become the industry standard small variant caller due to its high accuracy, and it has achieved top performance in a variety of public and third-party benchmarks [16,29,30,31]. DeepVariant and DNAscope are two newly developed variant callers based on machine learning methods [18,19,32,33]. It should be noted that both machine learning variant callers used in this study were optimized for the DNBSEQ™ platform through the use of in-house training data.

**Table 2.** Average variant calling performance of three replicates for PCR and PCR-free libraries using three variant callers. Variant calls from each library and with each variant caller were evaluated by Vcfeval in RTGtools against the NIST truth set at high-confidence regions. Average values from the same library construction method were generated and are shown here. “FL”represents full lane sequencing data, approximately 46x coverage.

| Pipeline | Variant type | Method | PCR |  |  | PCR-free |  |  |
| --- | --- | --- | --- | --- | --- | --- | --- | --- |
|  |  | Depth | 15x | 30x | FL | 15x | 30x | FL |
| GATK | SNPs | True positive | 3,106,478 | 3,193,586 | 3,198,933 | 3,126,614 | 3,199,331 | 3,202,461 |
|  |  | False positive | 9,615 | 2,806 | 2,479 | 8,544 | 2,586 | 2,347 |
|  |  | False negative | 103,778 | 16,671 | 11,324 | 83,643 | 10,926 | 7,796 |
|  |  | Precision | 99.69% | 99.91% | 99.92% | 99.73% | 99.92% | 99.93% |
|  |  | Sensitivity | 96.77% | 99.48% | 99.65% | 97.39% | 99.66% | 99.76% |
|  |  | F-score | 98.21% | 99.70% | 99.79% | 98.55% | 99.79% | 99.84% |
|  | InDels | True positive | 397,779 | 458,110 | 466,343 | 429,105 | 474,921 | 477,918 |
|  |  | False positive | 34,787 | 20,699 | 17,891 | 19,730 | 2,766 | 1,686 |
|  |  | False negative | 83,485 | 23,154 | 14,922 | 52,160 | 6,345 | 3,349 |
|  |  | Precision | 91.96% | 95.67% | 96.30% | 95.60% | 99.42% | 99.65% |
|  |  | Sensitivity | 82.65% | 95.19% | 96.90% | 89.16% | 98.68% | 99.31% |
|  |  | F-score | 87.05% | 95.43% | 96.60% | 92.27% | 99.05% | 99.48% |
| DeepVariant | SNPs | True positive | 3,174,776 | 3,205,991 | 3,207,021 | 3,184,757 | 3,207,148 | 3,207,769 |
|  |  | False positive | 12,952 | 2,913 | 1,871 | 7,837 | 2,111 | 1,691 |
|  |  | False negative | 35,481 | 4,266 | 3,236 | 25,499 | 3,109 | 2,488 |
|  |  | Precision | 99.59% | 99.91% | 99.94% | 99.75% | 99.93% | 99.95% |
|  |  | Sensitivity | 98.90% | 99.87% | 99.90% | 99.21% | 99.90% | 99.92% |
|  |  | F-score | 99.24% | 99.89% | 99.92% | 99.48% | 99.92% | 99.93% |
|  | InDels | True positive | 436,804 | 468,237 | 474,129 | 470,682 | 477,616 | 478,556 |
|  |  | False positive | 22,906 | 8,008 | 4,389 | 6,310 | 2,124 | 1,615 |
|  |  | False negative | 44,546 | 13,082 | 7,185 | 10,695 | 3,690 | 2,745 |
|  |  | Precision | 95.02% | 98.32% | 99.08% | 98.67% | 99.56% | 99.66% |
|  |  | Sensitivity | 90.74% | 97.28% | 98.51% | 97.78% | 99.23% | 99.43% |
|  |  | F-score | 92.83% | 97.80% | 98.80% | 98.23% | 99.39% | 99.55% |
| DNAScope | SNPs | True positive | 3,171,728 | 3,205,902 | 3,207,318 | 3,180,864 | 3,207,055 | 3,207,911 |
|  |  | False positive | 8,351 | 1,565 | 1,021 | 6,034 | 1,254 | 901 |
|  |  | False negative | 38,529 | 4,355 | 2,939 | 29,393 | 3,202 | 2,346 |
|  |  | Precision | 99.74% | 99.95% | 99.97% | 99.81% | 99.96% | 99.97% |
|  |  | Sensitivity | 98.80% | 99.86% | 99.91% | 99.08% | 99.90% | 99.93% |
|  |  | F-score | 99.27% | 99.91% | 99.94% | 99.45% | 99.93% | 99.95% |
|  | InDels | True positive | 443,656 | 466,964 | 471,037 | 466,852 | 477,969 | 479,103 |
|  |  | False positive | 21,653 | 8,632 | 5,773 | 5,617 | 1,620 | 1,202 |
|  |  | False negative | 37,609 | 14,301 | 10,229 | 14,413 | 3,299 | 2,164 |
|  |  | Precision | 95.35% | 98.19% | 98.79% | 98.81% | 99.66% | 99.75% |
|  |  | Sensitivity | 92.18% | 97.03% | 97.88% | 97.00% | 99.31% | 99.55% |
|  |  | F-score | 93.74% | 97.60% | 98.33% | 97.90% | 99.49% | 99.65% |

**Figure 3.**
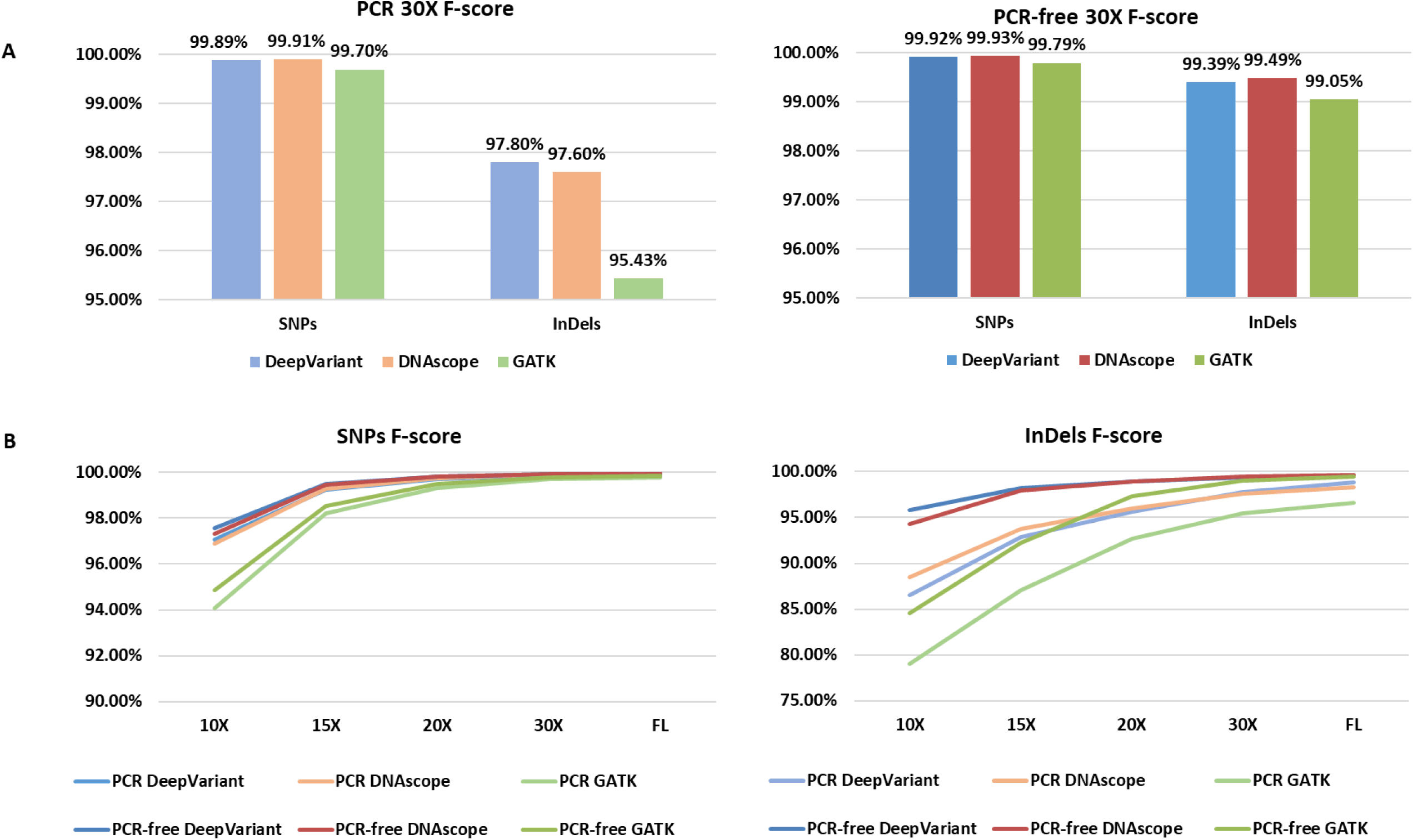
Variant calling performance in PCR vs. PCR-free libraries. (A) F-Scores of 30x sequencing PCR and PCR-free libraries were compared for accuracy performance using different variant callers. (B) Down-sampled datasets representing different sequencing depths were processed to examine the tolerance to shallow data from different variant callers. PCR-free libraries (dark color) and machine learning variant callers (blue and red) showed good accuracy at >15X that was superior to the PCR+GATK combination at 30X depth.

The variant calling matrix is highly reproducible in all three replicates for both PCR and PCR-free libraries (Supplementary Table 1S). Table 2 summarizes the average number of three replicates for different depths. At 30x depth, GATK called and marked as “Passed Filter” (named true positive, or TP for short) an average total of 3,651,696 TP variants for PCR libraries and 3,674,252 TP variants for PCR-free libraries, whereas DeepVariant and DNAscope showed higher sensitivity as they both detected additional TP variants for both PCR and PCR-free libraries. With all three callers, PCR-free libraries demonstrated a slight reduction in both the numbers of FP SNPs (from 2,806 to 2,586 in GATK, from 2,913 to 2,111 in DeepVariant, and from 1,565 to 1,254 in DNAscope) and the numbers of FN SNPs (from 16,671 to 10,926 in GATK, from 4,266 to 3,109 in DeepVariant, and from 4,355 to 3,202 in DNAscope) and a dramatic reduction in FP InDels (from 20,699 to 2,766 in GATK, from 8,008 to 2,124 in DeepVariant, and from 8,632 to 1,620 in DNAscope) and FN InDels (from 23,154 to 6,345 in GATK, from 13,082 to 3,690 in DeepVariant, and from 14,301 to 3,299 in DNAscope). This reduction in FPs and FNs leads to a slightly increased SNP F-score (harmonic mean of recall and precision) and a significantly increased InDel F-score for PCR-free libraries, suggesting more precise variant calling for all depths (Fig. 3a). As the highest accuracy combination, PCR-free data with DNAscope had the lowest FP SNPs, FP InDels, and FN InDels. As an additional evaluation, in selected “difficult” genome regions such as repeat regions and extreme GC regions, PCR-free libraries generally showed better InDel F-scores than PCR libraries and produce more faithful genome sequences for applications (Supplementary Fig. 2S).

To increase confidence in the performance of the PCR-free WGS workflow across a variety of sequencing depths, we generated additional 10x and 20x data points to conduct a 10x-46x low-depth test. As demonstrated in Fig. 3B, the reduction in coverage (i.e., 10x, 15x, and 20x) clearly affected the quality of variant calling from all methods. Nevertheless, the PCR-free method coupled with machine learning-based callers produced more accurate calling than pipelines involving PCR library construction or the GATK caller. Importantly, the SNP F-scores of DNAscope and DeepVariant for 15x PCR-free libraries were comparable to GATK for 30x PCR libraries (99.45%, 99.48%, and 99.70%, respectively), whereas the InDel F-scores of DNAscope and DeepVariant for 15x PCR-free libraries were significantly higher than that of GATK for 30x PCR libraries (97.90%, 98.23%, and 95.43%, respectively), indicating the potential to decrease sequencing cost while maintaining variant detection accuracy. Of note, DeepVariant showed the highest accuracy among all callers with 15x PCR-free libraries.

### Reproducibility of PCR-free libraries

We also conducted consistency analysis to determine the level of randomness introduced in library construction and sequencing and whether variant callers can help correct this randomness. The variant consistency of three replicates from PCR-free libraries was better than that observed with PCR libraries, especially for InDel consistency, and this trend is similar for all three variant callers (Fig. 4). This result is expected because the PCR step inevitably introduces random errors during amplification. The InDel consistency (represented by the portion of variants shared by all replicates) of PCR-free libraries was 84.2% with GATK, 86.5% with DeepVariant, and 89.1% with DNAscope (average 86.6%), which is approximately 20% greater than the InDel consistency seen with PCR libraries (63.2% with GATK, 66.9% with DeepVariant, and 68.9% with DNAscope (average 66.3%). We also observed that more than 99% of SNPs and approximately 98% of InDels in each library overlapped with at least one of the other two libraries. The consistency of SNPs among all three replicates was similarly high from both PCR and PCR-free libraries (three callers average 94.6% vs. 95.2%).

**Figure 4.**
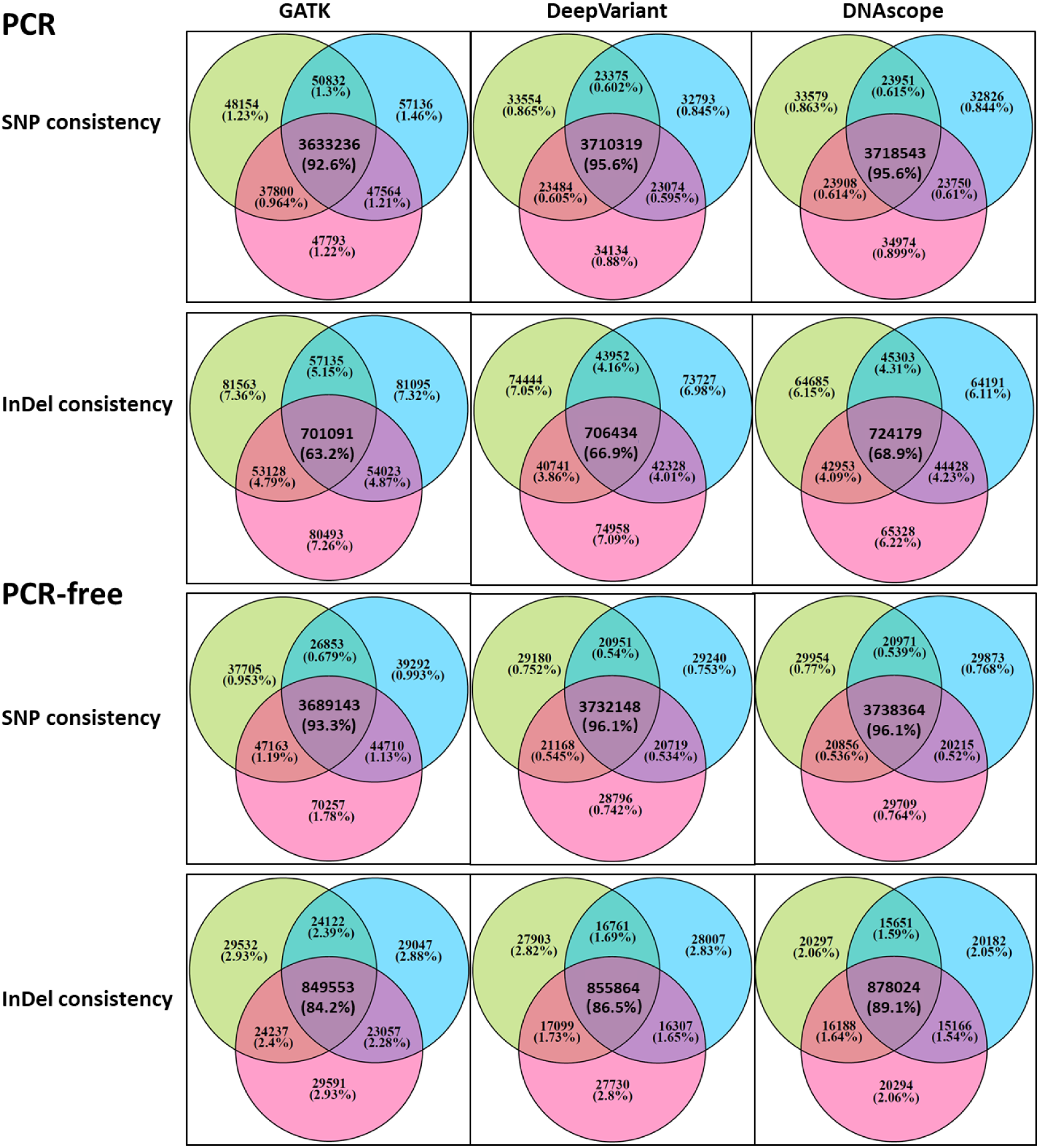
Variant consistency of 3 replicates of PCR and PCR-free libraries. Consistency analysis was conducted on the 3 libraries generated by the same library kit and variant calling pipelines. Venn diagrams were generated to show the common shared variants and the unique variants.

In contrast, we investigated the consensus calling on the same library among three callers. On average, the three pipelines for SNPs and InDels are 92.3% and 67.0% for PCR libraries and 92.6% and 82.7% for PCR-free libraries, indicating that PCR-free libraries generated more “clear” variant candidates that are less challenging for variant callers (Supplementary Fig. 3S).

### Evaluation of PCR-free WGS performance on different sequencing platforms

In addition to the library construction kit, the sequencing platform introduces additional bias or systematic variation that causes different performance. Here we compared two PCR-free libraries prepared using the MGIEasy PCR-Free DNA Library Pre Set and the MGIEasy FS PCR-Free DNA Library Pre Set and sequenced on MGISEQ-2000 with three datasets downloaded from the Illumina Basespace website to represent the performance of TruSeq PCR-free libraries sequenced on HiSeq4000, HiSeqXTen, or Novaseq platforms. The following three additional datasets were included in the comparison to provide further information: 1) library prepared with MGIEasy PCR-Free DNA Library Pre Set and sequenced on DNBSEQ-T7 [34]; 2) library prepared with MGIEasy PCR-Free DNA Library Pre Set with Illumina’s adapter and sequenced on Novaseq by a third-party sequencing service provider; 3) library prepared with research modifications of the MGIEasy kit and sequenced on MGISEQ-2000. It should be noted that libraries prepared with the MGIEasy kit used 1 μg or 250 ng DNA input, far less than the input for datasets downloaded from Illumina Basespace. All FASTQ files were processed in the same pipeline for read trimming/filtering, mapping, and variant calling using DNAscope (see Methods) to minimize bias or variation introduced at the secondary analysis stage.

From mapping the QC matrix, all three datasets generated from MGISEQ-2000 showed a significantly lower duplicate rate of approximately 1% or less, whereas all Illumina platform datasets, including the hybrid sample with MGI library prep, showed at least a 10% duplicate rate (Table 3). The “T7” dataset had a 3.60% duplication rate with 250 ng genomic DNA as input, which is still lower than Illumina’s duplicate rate. Thus, there was a more cost-effective generation of unique reads by DNBSEQ™. With DNAscope’s DNBSEQ™ and Illumina models applied separately, the SNP calling accuracy (represented by F-score) of all samples reached a similarly high level, except for Hiseq4000 (Table 3). For InDels, the two pure MGI pipeline (library prep + sequencing) - generated datasets and the “research library dataset” with 1µg as input all outperformed the three Illumina datasets with 2µg as input, showing less errors (FP and FN). This result is most likely due to the complete elimination of PCR during both library construction and sequencing procedures and confirms our hypothesis that PCR-generated DNA clusters frequently introduce clonal errors, especially InDels. Comparing the top performer from both sides—the “research library dataset” and the “TruSeq and Novaseq dataset”—the “research library dataset” had similar SNP calling but significantly lower false InDel calling with only 1,304 FPs, which is an ∼55% reduction from 2,879 FPs with the “TruSeq and Novaseq dataset”; 2,201 FNs in the “research library dataset” also represents an ∼48% reduction. The inclusion of data from the new DNBSEQ-T7 sequencer demostrates its accuracy and performance relative to other sequencing platforms. Although scale and cost has been prioritized in designing this ultra-high throughput sequencer, its sequencing accuracy has also been preserved, reaching the level of the “TruSeq and Novaseq dataset”.

**Table 3.** Mapping and Variant calling performance of PCR-free libraries sequenced on different sequencing platforms. DNBSEQ-T7 and MGISEQ-2000 data were generated according to PCR-free kit instructions or with research modifications (PCR-Free research lib).Illumina Hiseq4000, HiSeq xTen, and Novaseq data were downloaded from the Illumina Basespace website. To compare the sequence platform only, one PCR-free library was generated by MGIEasy FS PCR-Free DNA Library Prep Set using Illumina’s adapter and sequenced on Novaseq by a third-party sequencing service provider. Variants from each library were called using the DNAscope pipeline, and accuracy was evaluated by the vcfeval tool in RTGtools against the NIST truth set at high confidence regions.

| Library Kit |  | MGIEasy PCR-Free<br>DNA Library Prep<br>Set | MGIEasy FS PCR-Free DNA Library<br>Prep Set | MGIEasy PCR-Free<br>DNA Library Prep<br>Set | MGIEasy PCR-Free<br>DNA Library Prep<br>Set | PCR-Free<br>research lib | TruSeq DNA PCR-Free Library Prep Kits |  |  |
| --- | --- | --- | --- | --- | --- | --- | --- | --- | --- |
| DNA input |  | 250 ng | 1 µg | 1 µg | 1 µg | 1 µg | 2 µg | 2 µg | 2 µg |
| Sequencing platform |  | DNBSEQ-T7 | MGISEQ-2000 | MGISEQ-2000 | Novaseq | MGISEQ-2000 | Hiseq4000 | xTen | Novaseq |
| Average seq depth (X) |  | 30.46 | 30.68 | 30.91 | 30.75 | 30.6 | 30.02 | 31.5 | 30.43 |
| Mapping rate (%) |  | 99.99 | 99.89 | 99.86 | 99.82 | 99.83 | 99.03 | 99.85 | 99.94 |
| Duplicate rate (%) |  | 3.6 | 0.6 | 0.91 | 18.19 | 0.76 | 13.62 | 12.31 | 10.43 |
| Coverage at least 10X (%) |  | 98.64 | 98.73 | 98.69 | 98.83 | 98.82 | 98.8 | 98.91 | 98.87 |
| SNPs | True positive | 3,206,204 | 3,207,301 | 3,207,056 | 3,207,540 | 3,207,839 | 3,206,782 | 3,207,834 | 3,208,037 |
|  | False positive | 1,545 | 1,209 | 1,255 | 1,331 | 1,048 | 2,704 | 1,280 | 1,262 |
|  | False negative | 4,053 | 2,956 | 3,201 | 2,717 | 2,418 | 3,475 | 2,423 | 2,220 |
|  | Precision | 99.95% | 99.96% | 99.96% | 99.96% | 99.97% | 99.92% | 99.96% | 99.96% |
|  | Sensitivity | 99.87% | 99.91% | 99.90% | 99.92% | 99.92% | 99.89% | 99.92% | 99.93% |
|  | F-score | 99.91% | 99.94% | 99.93% | 99.94% | 99.95% | 99.90% | 99.94% | 99.95% |
| InDels | True positive | 476,673 | 478,047 | 478,004 | 477,283 | 479,066 | 469,353 | 477,420 | 477,017 |
|  | False positive | 2,266 | 1,882 | 1,595 | 2,479 | 1,304 | 7,642 | 2,598 | 2,879 |
|  | False negative | 4,592 | 3,218 | 3,265 | 3,984 | 2,201 | 11,913 | 3,846 | 4,250 |
|  | Precision | 99.53% | 99.61% | 99.67% | 99.48% | 99.73% | 98.40% | 99.46% | 99.40% |
|  | Sensitivity | 99.05% | 99.33% | 99.32% | 99.17% | 99.54% | 97.52% | 99.20% | 99.12% |
|  | F-score | 99.29% | 99.47% | 99.49% | 99.33% | 99.64% | 97.96% | 99.33% | 99.26% |

These data collectively highlighted that the DNBSEQ™ PCR-free WGS workflow provides clear benefits in sequencing efficiency and data quality based on parameters such as low duplication rate and high InDel calling accuracy.

## Discussion

The wide adoption of WGS can be credited to the dramatically decreasing cost of sequencing and to the potential downsides of WES, such as missing variants in the non-exome regions and inability to detect copy number variation. The development of machine-learning based secondary analysis tools also promotes the usage of WGS by shortening the analysis time from days to hours while attaining a higher variant calling accuracy.

### Advantages of PCR-free DNBSEQ™

In this study, we combined PCR-free-prepared WGS libraries with DNBSEQ™, thus excluding PCR from the entire sequencing process. This approach leads to a very low GC bias and overall coverage bias. We showed that PCR-free libraries had more accurate small variant (especially InDel) calls at 30x and other sequencing depths. This result confirmed a previous report that a PCR-free library had high-quality InDel calls [13]. This advantage expands when machine learning variant callers, such as DeepVariant and DNAscope, are applied instead of GATK. The fact that the InDel F-score of PCR-free libraries called by both machine learning callers at 15x depth is significantly better than that of PCR-prepared libraries called by GATK at 30x suggests that low-depth sequencing data could be used to conduct variant calling without compromising accuracy. Furthermore, true PCR-free WGS, enabled for the first time by high-quality PCR-free libraries and the PCR-free DNBSEQ™ platform, is expected to improve the detection of low-frequency somatic mutations.

The duplicate rate represents the proportion of duplicate reads from all the sequenced data. To ensure accuracy, the duplicated reads need to be removed for subsequent bioinformatics analysis. Therefore, for the same amount of raw data, a lower duplicate rate yields more usable clean data. The average duplicate rate of our PCR-free data on MGISEQ-2000 is approximately 1%, which is much lower than the duplicate rate for the PCR-free libraries (10.43%-13.62%) from the Illumina platforms. Thus, for 30x WGS, 90-100G raw bases is normally sufficient on MGISEQ-2000; however, approximately 20% more raw base (110-120G) is recommended on Illumina platforms. The superiority of the MGISEQ-2000 duplicate rate is due to the true PCR-free process, which employs no PCR in either library construction or sequencing workflow.

### Rapid WGS for infant genetic disorder diagnostics

Genetic disorders are among the top causes of morbidity and mortality in infants; the newly developed rapid whole-genome sequencing (rWGS) shows the power to diagnose genetic disorders quickly, thus enabling healthcare providers to generate or change their management plan accordingly and thus improve outcomes for acutely ill infants [35]. To facilitate this clinical utility, it is essential that the whole process, from sample collection to diagnostic report review and signing, can be completed within 2 days. As a result, there is urgent demand to decrease the turn-around time of library construction, sequencing, and secondary analysis to 24 hours, thus providing more time for candidate variant clinical annotation and board review.

Compared to traditional PCR-inclusive library construction methods, the PCR-free method skips amplification and therefore significantly reduces the turn-around time. Moreover, the enzymatic shearing method makes this method more amenable to automation and further decreases hands-on time in library construction. The new DNBSEQ-T7 sequencer, with super high-throughput of up to 6T per run, can finish within 24 h, which greatly decreases the sequencing time compared with MGISEQ-2000 and provides the possibility to realize rWGS. For secondary analysis, the most time-consuming steps are read alignment and variant calling. The traditional BWA GATK pipeline takes more computational time than is optimal, but some alternative pipelines have been proposed and developed to meet the accelerated speed requirements, including CPU-based tools, such as Strelka2 [36], DNAseq [37], and SpeedSeq [38]; FPGA&CPU-based instruments, such as DRAGEN [39] and MegaBOLT [40]; and the GPU/TPU implemented Deepvariant [18]. All these advanced bioinformatics tools are compatible with data generated from MGI’s library construction kit and MGI sequencing platforms.

### Structural Variation (SV) and Copy Number Variation (CNV)

SV and CNV are two critical clinical parameters for which researchers choose WGS instead of WES. Although not designed as the primary goal for this benchmark study, the data generated allow us to investigate whether our PCR-free method improves SV and CNV detection accuracy. SV was called by DNAscope in default parameters, and CNV was called by the GATK 4.1.2 pipeline. Supplementary Fig.4S demonstrates that the PCR-free method likely helps improve both the sensitivity and specificity of SV calling compared to PCR-based library construction methods, because more SV events were called in the PCR-free group and they reached higher consistency among three replicate samples. A key point in detecting SV events is correctly detecting breakpoints, which relies on sufficient coverage across targets and less errors that generate false positives. Obviously, PCR-free libraries will benefit this detection.

Germline CNV for all six testing samples were called by GATK 4.1.2, and approximately 2,500 CNV events (mainly deletions) were identified from each sample. When conducting 3-way comparison to analyze the reproducibility among replicates, the PCR-free group showed a slightly higher number of common CNV events, but the overall difference compared to the PCR group was negligible (Supplementary Fig. 5S).

### Clinical utility of higher WGS accuracy

Clinical WGS has begun to show its potential in rare disease diagnostic capacity, because WGS can quickly cover the whole genome and identify clinically meaningful variants, especially in UTRs or promoter regions that panel sequencing or WES would fail to detect. The increased variant detection accuracy using PCR-free library construction and machine learning-based variant calling pipelines clearly increased WGS variant calling accuracy and will therefore surely add value to diagnostic applications. In other words, for regions that PCR-based WGS fails to generate sufficient read coverage or consistently generates wrong variant calling, PCR-free WGS will be able to provide correct SNP/InDel information. From the six DNBSEQ™ datasets and three Illumina datasets evaluated in this study, we found two example genes for which all three PCR-free libraries showed accurate variant calling in the gene-coding or UTR regions but for which all three PCR-based libraries generated FNs. These two genes were ATK1 and GNAS (Supplementary Fig. 6S, Supplementary Fig. 7S). Similarly, we also showed one example gene, MAF (Supplementary Fig. 8S), for which the DNBSEQ™ PCR-free method really excelled; all three Illumina datasets and both MGI PCR datasets failed to detect a C to CT insertion in this gene. These three genes code clinically meaningful proteins in which failed variant detection could lead to mis-diagnosis. For example, ATK1 (ATK serine / threonine kinase 1) is associated with multiple clinical phenotypes, including breast cancer (MIM #114480), colorectal cancer (MIM #114500), Cowden syndrome 6 (MIM #615109), and ovarian cancer (MIM #167000). The T to TC insertion at locus Chr14:105262025 and the CG to GC SNP at Chr14:105262041 may cause malfunction and introduce a disease phenotype, and only PCR-free libraries were able to identify these critical variants.

### Future improvements

As a benchmark project, this study shows the current performance of tools and pipelines used in library preparation, sequencing, and data analysis. MGI’s PCR-free WGS sets provide solutions for both ultrasonic and enzymatic genomic DNA fragmentation and demonstrate good reproducibility from a broad range of DNA input (200 ng - 1 µg for MGIEasy PCR-Free DNA Library Prep Set and 50 ng - 1 µg for MGIEasy FS PCR-Free DNA Library Prep Set). Modified protocols based on these sets can yield even better InDel accuracy, which indicates that room exists to upgrade the PCR-free WGS sets. As expected, emerging technologies will continually push the upper limit of sequencing accuracy. For example, the performance of new DNBSEQ-T7 sequencer [34] was briefly displayed in this study. Among other sequencers in development, this machine can decrease sequencing cost per genome or provide deeper coverage and further shorten the sequencing time as needed. The poor capacity for structural variation detection due to the nature of short-read sequencing can be greatly compensated by applying long fragment read (LFR) barcoding technologies [41,42,43]. For some clinical samples such as cfDNA, it is impractical to obtain 200 ng for library preparation. With the developed method based on MGI PCR-Free sets and a pooling sequencing strategy, we successfully generated good data from 10 ng cfDNA in multiple studies (data not shown).

On the analysis side, the current GATK best practice pipeline was developed and tuned based on Illumina data and does not officially support DNBSEQ™ -generated data as of December 2018 according to the GATK team’s response in forum [44]. Certain error correction steps in the pipeline, such as BQSR and VQST, were not developed for DNBSEQ™ data and thus may generate non-optimized results compared to Illumina sequencer-generated data. This view is supported by two recently published benchmark studies [45,46]; DNBSEQ™ data analyzed by Strelka2 and DeepVariant showed comparable accuracy with Illumina data, but DNBSEQ™ data analyzed by GATK returned a worse accuracy.

Both DeepVariant and DNAscope rely on proper model training, which requires sufficient sample numbers prepared using the same library construction method and sequencing platform as the testing samples. In this study, we do not believe this requirement was fully met, especially for DNAscope. For example, only 30x and higher depth datasets were included into the training set, but testing on 15x depth data was conducted. The negative effect of DNAscope lacking a proper 15x depth training dataset is shown when comparing its 10x and 15x accuracy to that of DeepVariant, for which the training dataset included 15x depth data. Another point worth noticing is that using a single model for all library kits/assays and sequencers could sacrifice accuracy for individual cases. It is best for users to train individual models for each individual case (i.e., the combination of sample prep kit, library construction kit, and sequencing platform) to achieve optimal variant calling accuracy. With all the above-mentioned improvements for future WGS cohort studies, the unprecedented data generation speed and quality will help to answer difficult genetic questions and move the genomics field into a new era of broad clinical use.

## Conclusions

In this study, we present an advanced WGS solution using an entirely PCR-free MPS workflow—DNBSEQ™ PCR-free WGS, which the whole process is not only PCR-free during library preparation but also PCR-free during sequencing. Data from repeatability tests show that the DNBSEQ™ PCR-free WGS has low coverage bias, good repeatability and consistency, and the minimum starting amount of genomic DNA can be low as 50ng. Analysis results of DNBSEQ™ PCR-free comparing with DNBSEQ™ PCR and illumina’s PCR-free demonstrate that DNBSEQ™ PCR-free have the minimum systematic errors,which provides clear benefits in sequencing efficiency and data quality based on parameters such as low duplication rate and high InDel calling accuracy. The MGI’s PCR-free toolkit (including ultrasonic or enzymatic DNA fragmentation MGI library preparation kits and DNBSEQ™ series sequencer) combinded with ML-based variant calling pipelines (DeepVariant or DNAscope) can achieve even better data quality in terms of SNP and InDel F-scores,moreover,this combination also provide a possibility for clinical diagnostics using cost-efficient middle-pass WGS (i.e.,15X,10X WGS). We believe that DNBSEQ™ PCR-free WGS is a powerful solution for genome research, cohort studies and precision medicine.Further optimizations in quality, speed and cost throughout the entire MPS from library construction to data analysis will help the universal application of DNBSEQ™ PCR-free WGS.

## Methods

### DNA preparation

Genomic DNA from NA12878 cells (RRID:CVCL_7526) was purchased from the Coriell Institute. Genomic DNA was quantified using a Qubit™ 3 Fluorometer (Life Technologies, Carlsbad, CA, USA). The size and quality of genomic DNA were confirmed by running 0.8% agarose gels.

### PCR-free library preparation

Ultrasonically fragmented PCR-free libraries and enzymatically fragmented PCR-free libraries were constructed using the MGIEasy PCR-Free DNA Library Prep Set (MGI, cat. no. 1000013453) and the MGIEasy FS PCR-Free DNA Library Prep Set (MGI, cat. No. 1000013455), respectively.

For ultrasonically fragmented PCR-free libraries, genomic DNA was fragmented to 100-600 bp with peak size of 350-400 bp using an LE220 Ultrasonicator (Covaris, Woburn, MA, USA). For FS PCR-free libraries, genomic DNA was fragmented to 100-1000 bp with peak size of 350-475 bp using an enzymatic shearing method. Subsequently, fragmented DNA with a size range of 200-450 bp was selected using MGIEasy DNA Clean Beads (MGI, cat. no. 1000005279) and attached with DNBSEQ™ adapters according to kit instructions. We also followed the protocols from the kit to prepare single-stranded DNA (ssDNA) circles and quantified these on a Qubit™ 3 Fluorometer.

The library preparation procedure for research libraries was similar to that used with the MGIEasy PCR-Free DNA Library Prep Set, except for the size selection and single strand degeneration methods.

### PCR library preparation

PCR libraries were prepared using the same procedure of DNA fragmentation, end repair, and adapter ligation with the MGIEasy PCR-Free DNA Library Prep Set (Cat. no. 1000013453) as described above. After adapter ligation, the reaction product was purified with MGIEasy DNA Clean Beads (MGI, cat. no. 1000005279), and the ligation products were subjected to PCR amplification following instructions from the KAPA HiFi HotStart ReadyMix (KAPA BIOSYSTEMS, KK2602). A total of 6 cycles (95°C 3 min; 6 cycles of 98°C 20 s, 60°C 15 s, and 72°C 60 s; 72°C 10 min; 4°C forever) were performed in a volume of 100 µl. After bead purification using MGIEasy DNA Clean Beads and quantification using a Qubit™ 3 Fluorometer, 1 pmol PCR product was used for single strand molecule circularization according to the ssCir formation protocol from the MGIEasy kit.

### Sequencing

Whole genome sequencing was performed on the DNBSEQ™ platforms MGISEQ-2000 using paired-end 150-bp (PE150) reads and on DNBSEQ-T7 using PE150 reads. Before sequencing, 75 fmol ssDNA from PCR-free libraries or 40 fmol single-strand circle DNA from PCR libraries were used to prepare DNA nanoballs (DNBs) according to the kit instructions from the MGISEQ-2000RS High-throughput Sequencing Set (FCL PE150) (MGI, cat. no. 1000012555). Seventy-five fmol ssDNA from PCR-free libraries was used to prepare DNBs according to kit instructions from the DNBSEQ-T7RS High-throughput Sequencing Set (PE150) (MGI, cat. no. 1000016106). Subsequently, DNBs were loaded onto the sequencing slide, and PE150 sequencing was conducted on DNBSEQ™ platforms using MGISEQ-2000RS or DNBSEQ-T7RS High-throughput Sequencing Sets.

### GC bias analysis

To explore GC bias, we sequenced two bacteria samples (Table 2S) on MGISEQ-2000 using PE150 reads. Filtered reads were aligned to reference genomes by Burrows-Wheeler aligner (BWA) [47]. To investigate the relationship between GC bias and read coverage, we scanned the genome with a sliding window of default size (100 bases). GC content and average read coverage were calculated within each window. Read coverage was normalized to the mean value such that the results would not scale with the total amount of data. In addition, we eliminated the data points whose coverage was more than twice the mean coverage because they likely represented repeats. Finally, we fit the remaining data points by a straight line and defined the slope as the degree of GC bias in the real data.

### Read filtering and mapping

As the first step, raw reads sequenced from PCR or PCR-free libraries were debarcoded by Seqtk [48] with default parameters. Split reads were then pre-processed by SOAPnuke to generate filtered reads [49]. During this filtering process, reads containing more than 10% of ‘N’ or 50% of the base quality score lower than 12 were removed. Adapters were trimmed off, and then alignment of all reads against the human reference genome with decoy sequencing hs37d5 (or reference genome sequences of two bacteria for GC bias analysis) was performed using BWA with default parameters. The output SAM file was converted to a BAM file and sorted by Samtools [50]. Lastly, duplicates were marked by Picard [51] to prepare both BAM files for variant calling by GATK, DeepVariant, and DNAscope.

### Running GATK

SNP and InDel calling was performed according to GATK (version 3.3) best practice [52]. Reads surrounding InDels were re-aligned, and base quality scores were recalibrated. HaplotypeCaller was used to call variants in gVCF mode on each chromosome. Genotyping on the gVCF files was performed using GenotypeGVCFs with parameters as follows: -stand_call_conf 30 and -stand_emit_conf 10. SNPs and InDels were separated using SelectVariants tool. Variant quality score recalibration (VQSR) was performed to filter low-quality variants. SNP annotation --ts_filter_level was used for calculation and filtered at a 99.9% level, whereas for InDels, --ts_filter_level was used for calculation and filtered at a 99.9% level of the true sensitivity.

### Running DeepVariant

Taking advantage of the state-of-the-art deep-learning technique for image classification, DeepVariant (V0.7.0 in this study) can achieve a higher accuracy for bioinformatics analysis. The genome in a bottle (GIAB) truth set and corresponding fastq reads were utilized as a training dataset to train a convolutional neural network (CNN) model. As an alternative to GATK HaplotypeCaller, DeepVariant accepts aligned reads (e.g., BAM files) as input. In DeepVariant, candidate variants are carefully filtered along the genome and classified into three genotype states, homozygous reference (hom-ref), heterozygous (het), or homozygous alternate (hom-alt) according to the previously trained CNN model.

To achieve best calling performance, we fine-tuned the CNN model in DeepVariant using a set of PCR-free data, including 30x and 15x DNBSEQ™ PCR-free sequencing data of HG001 and HG005 samples. The fine-tuned model is accessible at [53].

### Running DNAscope

Sentieon DNAscope (versions 201808.01 and 201808.05 were used in this study) uniquely combines the well-established methods from haplotype-based variant callers with machine learning to achieve improved accuracy. As a successor to GATK HaplotypeCaller, DNAscope uses an identical logical architecture of active region detection, local haplotype assembly, and read-likelihood calculation (Pair-HMM) to produce variant candidates. DNAscope produces additional informative variant annotations used by the machine learning model. Annotated variant candidates are then passed to a machine learning model for variant genotyping, resulting in improvements in both variant calling and genotyping accuracy.

For this study, DNBSEQ™ model for DNAscope was constructed using publicly available data from the HG001 and HG005 GIAB samples downloaded from the NIST GIAB FTP site along with proprietary 30x HG001 samples. The Illumina model for DNAscope was also trained using a subset of the GIAB HG001 and HG005 data. None of the tested samples were used during model training. Training was performed across all chromosomes with the exception of chromosome 20.

### Variant accuracy evaluation

All VCF files generated from this benchmark study were collected for accuracy evaluation. First, they were separated into SNP and InDel subgroups, and each subgroup was then compared against the NIST truth set using Vcfeval from RTGtools [54] to calculate an F-score as a representation of accuracy.

## List of abbreviations

BWA: Burrows-Wheeler aligner
CNN model: convolutional neural network model
CNV: Copy Number Variation
DNBs: DNA nanoballs
FP: false positive
FN: false negative
GATK: Genome Analysis Toolkit
GIAB: genome in a bottle
hom-ref: homozygous reference
het: heterozygous
hom-alt: homozygous alternate
InDels: insertions and deletions
LFR: long fragment read
MPS: massively parallel sequencing
ML: machine learning
NGS: next-generation sequencing
PCR: Polymerase chain reaction
Pair-HMM: read-likelihood calculation
rWGS: rapid whole-genome sequencing
SNPs: single-nucleotide polymorphisms
ssDNA: single-stranded DNA
SPRI: solid-phase reversible immobilization
SV: Structural Variation
VQSR: Variant quality score recalibration
WGS: whole genome sequencing
WES: whole-exome sequencing

## Declarations

### Ethics approval and consent to participate

The Institutional Review Board on Bioethics and Bio-safety of BGI (BGI-IRB), NO.FT18152 has approved this study.

### Availability of data and materials

The data reported in this study are available in the CNGB Nucleotide Sequence Archive (https://db.cngb.org/cnsa; accession number CNP0000602, CNP0000466).

All other data used here are included within the published article and additional files.

### Authors’ contributions

Conception and design of study: XZ, YJ,RD;

Acquisition of data: HS, PL, YX, QL, XW,LW, TB, AA,YL, MG, JL, NB, ZM, HR;

Analysis and/or interpretation of data: HS, ZL, SS, XL, Yinlong X, WT, Zhe L,XZ, YJ,RD;

Drafting the manuscript: HS, PL, ZL, SS, YX, JZ, LW, Zhe L, XZ, YJ,RD;

Revising the manuscript: XL, Yinlong X, SD, XZ,YJ,RD;

Supervision and resource support: FC, HJ, CX, SD, AC, WZ, FM, XX;

Fund acquisition: FC, RD;

Approval of the version of the manuscript to be published: All the authors listed in the title page.

## Acknowledgements

We would like to acknowledge the ongoing contributions and support of all MGI Tech Co., Ltd., BGI-Shenzhen and Complete Genomics Inc. employees, and we also acknowledge Frank Hu from Sentieon for commenting on the manuscript.

