## Supplementary Material for "Advanced Whole Genome Sequencing Using an Entirely PCR-free Massively Parallel Sequencing Workflow"

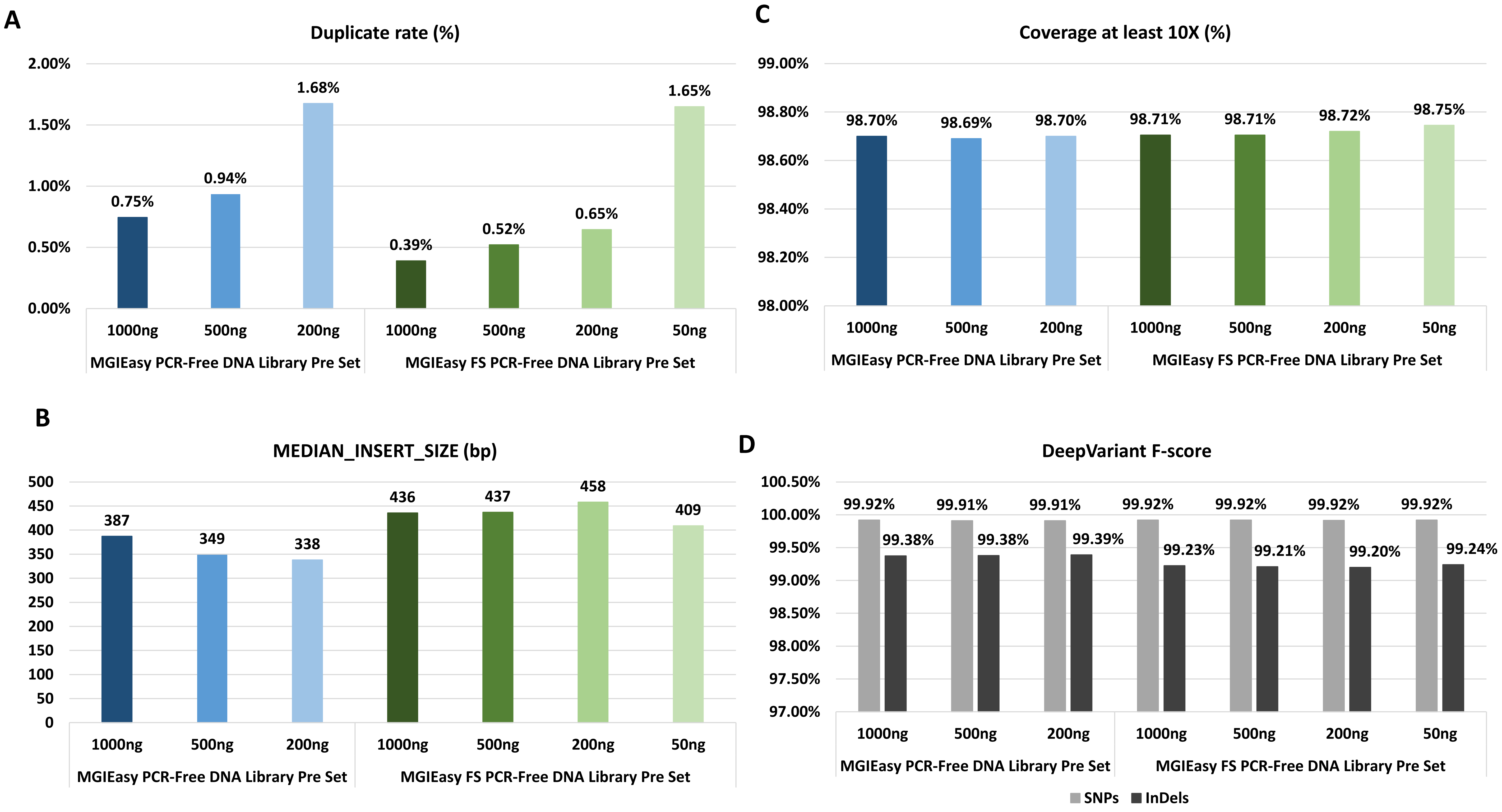


**Supplementary Figure 1S. Data performance with different library inputs using either ultrasonic shearing (MGIEasy PCR-Free DNA Library Prep Set) or enzymatic shearing (MGIEasy FS PCR-Free DNA Library Prep Set).** Each condition was repeated twice, and the average value is displayed.

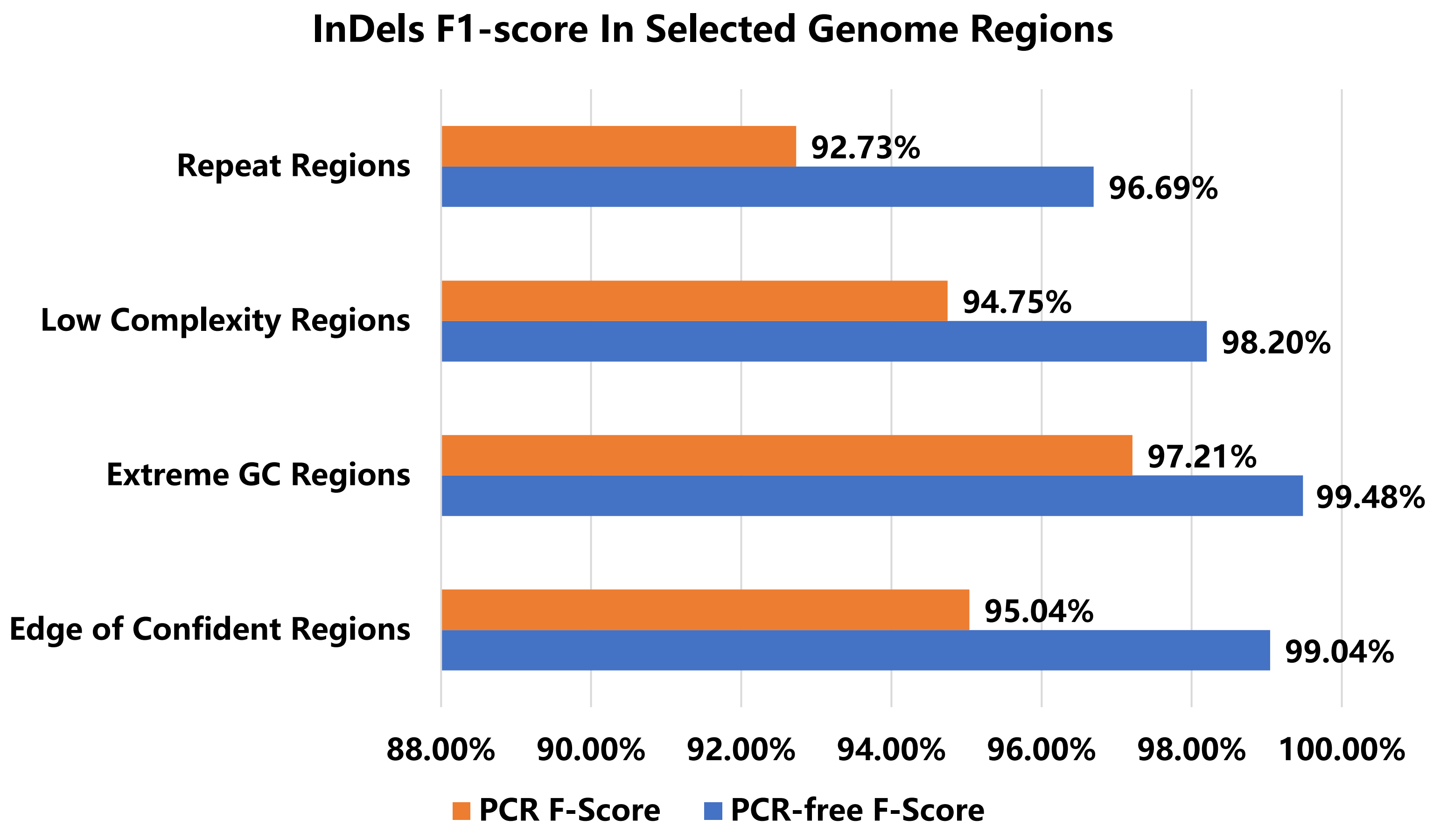


**Supplementary Figure 2S. InDels F-score in Selected Genome Regions.** In comparison between one PCR sample and PCR-free sample, a number of genome subsets regions were selected, and F1-scores generated from Sentieon DNAscope were compared within each region. InDels showed more significant differences between two samples, and especially in the 4 categories above. These categories are selected from GA4GH stratification regions with category names changed for easier understanding. Repeat Regions matches category“lowcmp_AllRepeats_51to200bp_gt95identity_merged”; Low Complexity Regions matches category “lowcmp_Human_Full_Genome_TRDB_hg19_150331_all_merged”; Extreme GC Regions matches category “gclt30orgt55”; Edge of Confident Regions matches category "TS_boundary”.

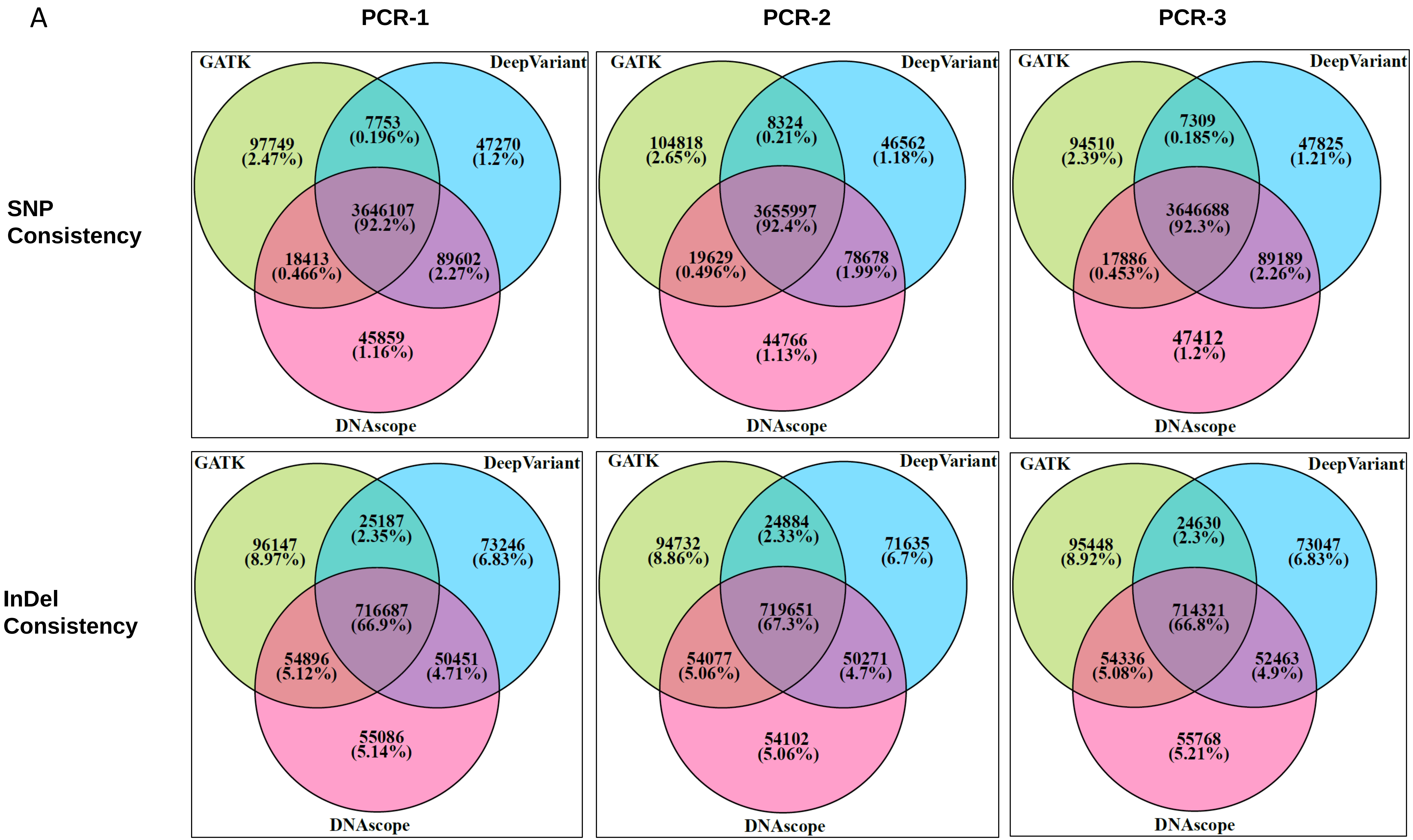


**Supplementary Figure 3S. Consistency of three pipelines for SNPs and InDels.** (A) Consistency analysis was conducted on the 3 variant calling pipelines on the same PCR library. Venn diagrams were generated to show the common shared variants and the unique variants.

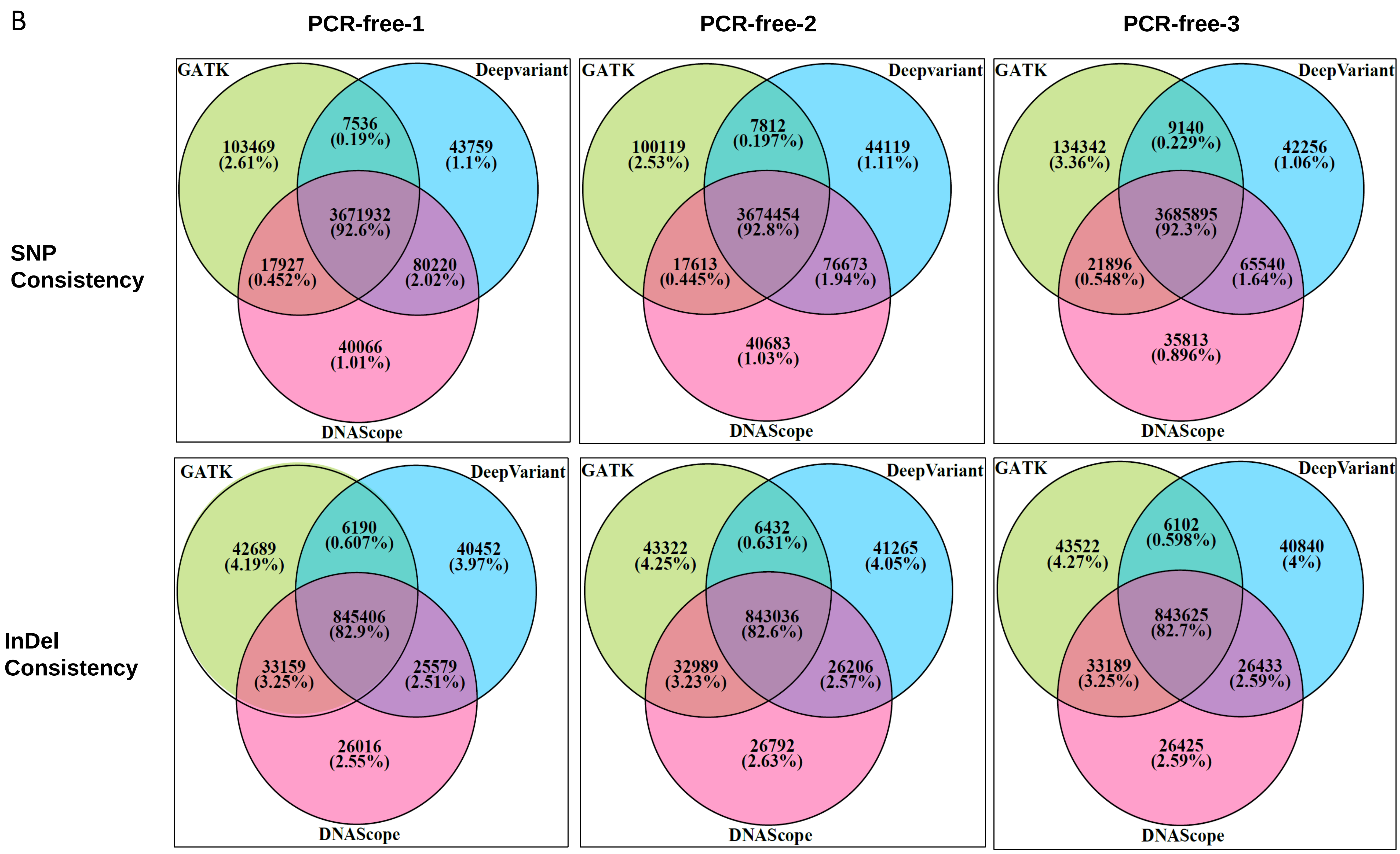


**Supplementary Figure 3S. Consistency of three pipelines for SNPs and InDels.** (B) Consistency analysis was conducted on the 3 variant calling pipelines on the same PCR-free library. Venn diagrams were generated to show the common shared variants and the unique variants.

A

| Samples | TRA | DEL | DUP | INV | INS |
|---|---|---|---|---|---|
| PCR-1_30X | 203 | 3,528 | 175 | 227 | 99 |
| PCR-2_30X | 205 | 3,488 | 138 | 224 | 99 |
| PCR-3_30X | 204 | 3,477 | 172 | 225 | 95 |
| PCR-free-1_30X | 293 | 3,684 | 189 | 231 | 104 |
| PCR-free-2_30X | 288 | 3,625 | 182 | 254 | 106 |
| PCR-free-3_30X | 251 | 3,689 | 168 | 256 | 123 |

B

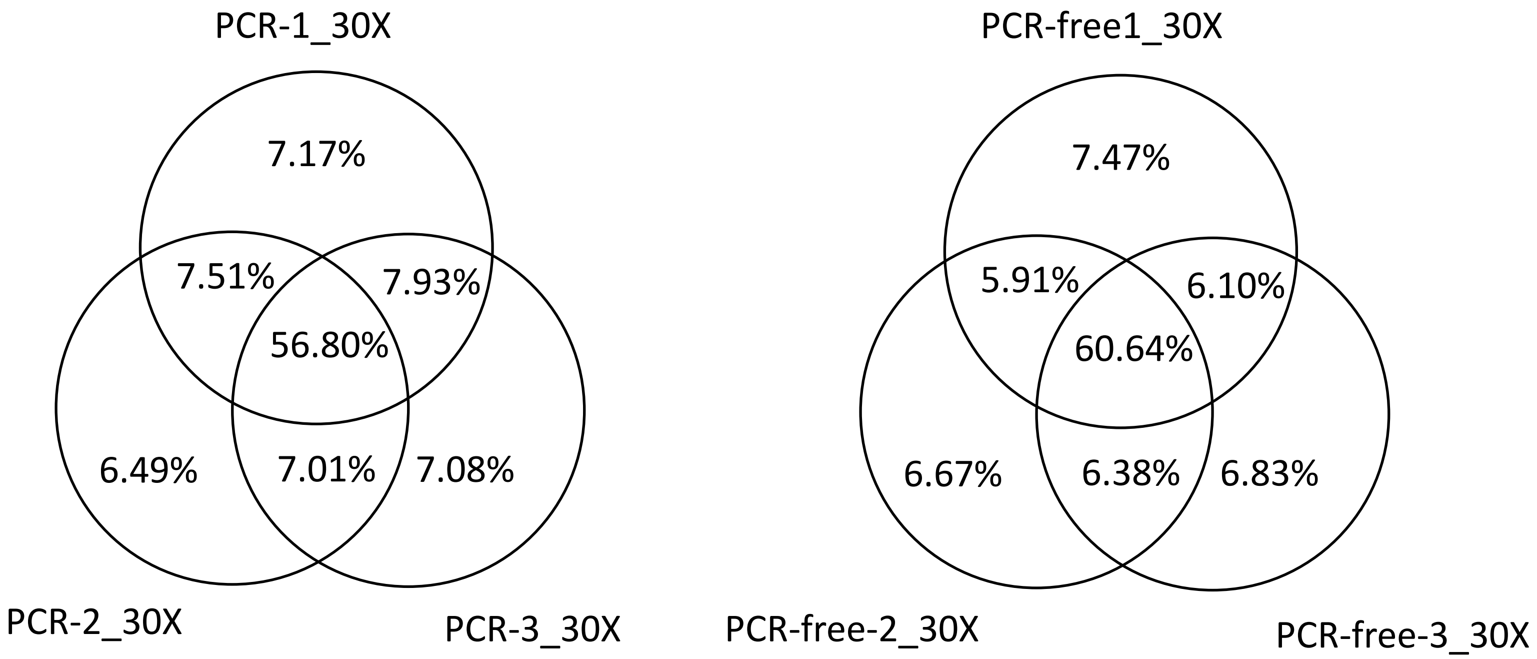


**Supplementary Figure 4S**. (A) SV events detected by DNAscope in 6 testing samples. SV events were reported into one of the five categories Translocation, Deletion, Duplication, Inversion, and Insertion. PCR-free libraries showed higher numbers of reported SV events and (B) higher consistency among all 3 samples.

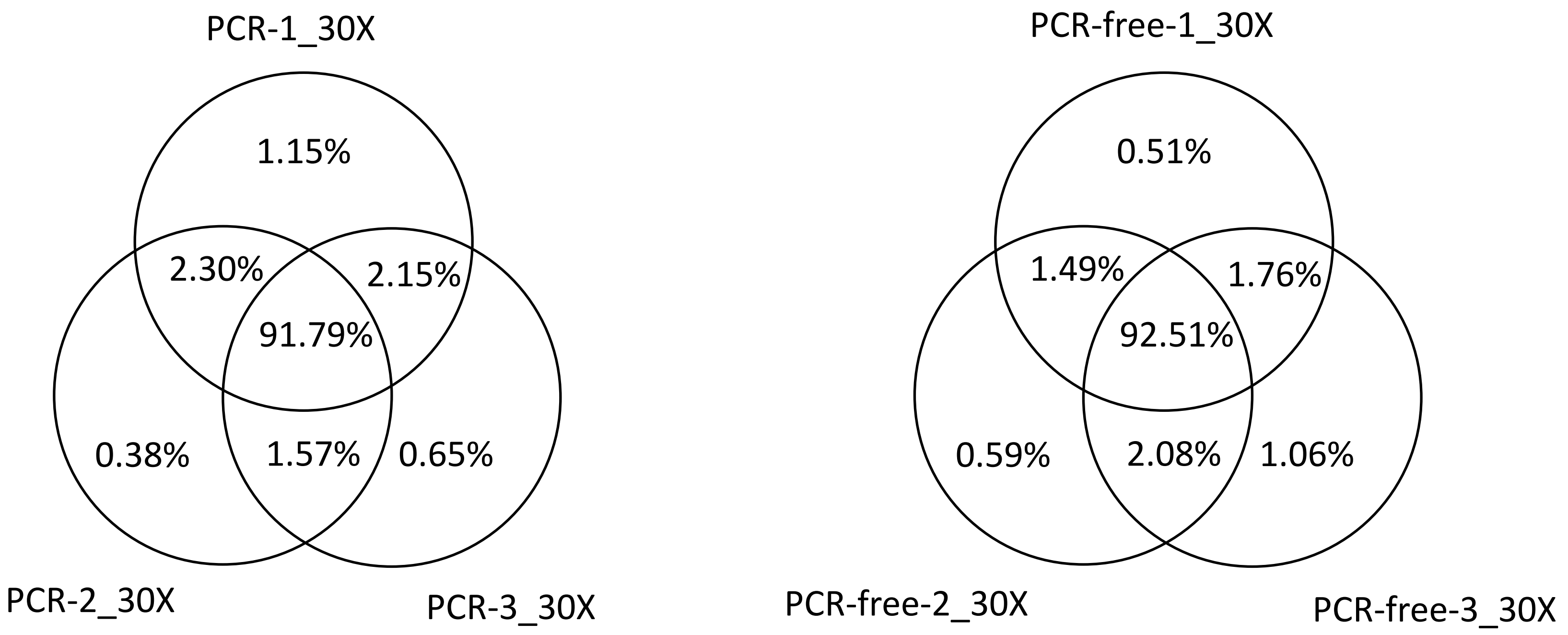


**Supplementary Figure 5S**. CNV events detected by DNAscope in 6 testing samples. 3-way comparison was conducted to assess consistency among replicates.

| Pipeline | Variant type | Method | PCR-1 | | | PCR-2 | | | PCR-3 | | | PCR-free-1 | | | PCR-free-2 | | | PCR-free-3 | | |
|---|---|---|---|---|---|---|---|---|---|---|---|---|---|---|---|---|---|---|---|---|
| | | Depth | 15x | 30x | FL | 15x | 30x | FL | 15x | 30x | FL | 15x | 30x | FL | 15x | 30x | FL | 15x | 30x | FL |
| | | True positive | 3110618 | 3193126 | 3200655 | 3108081 | 3195584 | 3197789 | 3100734 | 3192049 | 3198355 | 3128229 | 3198582 | 3201565 | 3123297 | 3198921 | 3202891 | 3128317 | 3200489 | 3202928 |
| | | False positive | 9634 | 2770 | 2750 | 9352 | 2939 | 2205 | 9858 | 2708 | 2483 | 8708 | 2403 | 2570 | 8559 | 2268 | 2385 | 8364 | 3086 | 2087 |
| | | False negative | 99638 | 17131 | 9602 | 102175 | 14673 | 12468 | 109522 | 18208 | 11902 | 82028 | 11675 | 8692 | 86960 | 11336 | 7366 | 81940 | 9768 | 7329 |
| | SNPs | Precision | 99.69% | 99.91% | 99.91% | 99.70% | 99.91% | 99.93% | 99.68% | 99.92% | 99.92% | 99.72% | 99.92% | 99.92% | 99.73% | 99.93% | 99.93% | 99.73% | 99.90% | 99.93% |
| | | Sensitivity | 96.90% | 99.47% | 99.70% | 96.82% | 99.54% | 99.61% | 96.59% | 99.43% | 99.63% | 97.44% | 99.64% | 99.73% | 97.29% | 99.65% | 99.77% | 97.45% | 99.70% | 99.77% |
| | | F-score | 98.27% | 99.69% | 99.81% | 98.24% | 99.73% | 99.77% | 98.11% | 99.67% | 99.78% | 98.57% | 99.78% | 99.82% | 98.49% | 99.79% | 99.85% | 98.58% | 99.80% | 99.85% |
| GATK | | | | | | | | | | | | | | | | | | | | |
| | | True positive | 398324 | 458096 | 466055 | 398912 | 458728 | 466804 | 396100 | 457507 | 466171 | 429199 | 475007 | 477924 | 428632 | 474850 | 477933 | 429485 | 474906 | 477897 |
| | | False positive | 35000 | 21111 | 18550 | 34291 | 20094 | 17163 | 35069 | 20892 | 17960 | 19737 | 2741 | 1688 | 19793 | 2775 | 1719 | 19661 | 2782 | 1650 |
| | | False negative | 82941 | 23168 | 15210 | 82352 | 22536 | 14462 | 85162 | 23757 | 15094 | 52066 | 6258 | 3343 | 52632 | 6417 | 3334 | 51783 | 6361 | 3370 |
| | InDels | Precision | 91.92% | 95.59% | 96.17% | 92.08% | 95.80% | 96.45% | 91.87% | 95.63% | 96.29% | 95.60% | 99.43% | 99.65% | 95.59% | 99.42% | 99.64% | 95.62% | 99.42% | 99.66% |
| | | Sensitivity | 82.77% | 95.19% | 96.84% | 82.89% | 95.32% | 96.99% | 82.30% | 95.06% | 96.86% | 89.18% | 98.70% | 99.31% | 89.06% | 98.67% | 99.31% | 89.24% | 98.68% | 99.30% |
| | | F-score | 87.10% | 95.39% | 96.50% | 87.24% | 95.56% | 96.72% | 86.82% | 95.35% | 96.58% | 92.28% | 99.06% | 99.48% | 92.21% | 99.04% | 99.47% | 92.32% | 99.05% | 99.48% |
| | | True positive | 3175227 | 3206052 | 3207074 | 3174871 | 3205898 | 3206874 | 3174231 | 3206023 | 3207115 | 3185071 | 3207185 | 3207765 | 3184356 | 3207120 | 3207784 | 3184844 | 3207140 | 3207757 |
| | | False positive | 12381 | 2879 | 1875 | 13175 | 2917 | 1886 | 13301 | 2942 | 1851 | 7973 | 2119 | 1700 | 7830 | 2138 | 1698 | 7709 | 2077 | 1675 |
| | | False negative | 35030 | 4205 | 3182 | 35386 | 4359 | 3383 | 36026 | 4234 | 3142 | 25185 | 3072 | 2492 | 25900 | 3137 | 2473 | 25413 | 3117 | 2500 |
| | SNPs | Precision | 99.61% | 99.91% | 99.94% | 99.59% | 99.91% | 99.94% | 99.58% | 99.91% | 99.94% | 99.75% | 99.93% | 99.95% | 99.75% | 99.93% | 99.95% | 99.76% | 99.94% | 99.95% |
| | | Sensitivity | 98.91% | 99.87% | 99.90% | 98.90% | 99.86% | 99.89% | 98.88% | 99.87% | 99.90% | 99.22% | 99.90% | 99.92% | 99.19% | 99.90% | 99.92% | 99.21% | 99.90% | 99.92% |
| | | F-score | 99.26% | 99.89% | 99.92% | 99.24% | 99.89% | 99.92% | 99.23% | 99.89% | 99.92% | 99.48% | 99.92% | 99.93% | 99.47% | 99.92% | 99.94% | 99.48% | 99.92% | 99.93% |
| DeepVariant | | | | | | | | | | | | | | | | | | | | |
| | | True positive | 436833 | 468201 | 474042 | 437513 | 468409 | 474219 | 436066 | 468101 | 474127 | 470823 | 477670 | 478584 | 470466 | 477597 | 478546 | 470756 | 477582 | 478538 |
| | | False positive | 23254 | 8085 | 4458 | 22359 | 7786 | 4257 | 23105 | 8152 | 4453 | 6233 | 2111 | 1614 | 6514 | 2120 | 1614 | 6182 | 2141 | 1616 |
| | | False negative | 44515 | 13120 | 7283 | 43834 | 12915 | 7093 | 45290 | 13212 | 7179 | 10553 | 3641 | 2719 | 10915 | 3711 | 2756 | 10617 | 3718 | 2760 |
| | InDels | Precision | 94.95% | 98.30% | 99.07% | 95.14% | 98.36% | 99.11% | 94.97% | 98.29% | 99.07% | 98.69% | 99.56% | 99.66% | 98.63% | 99.56% | 99.66% | 98.70% | 99.55% | 99.66% |
| | | Sensitivity | 90.75% | 97.27% | 98.49% | 90.89% | 97.32% | 98.53% | 90.59% | 97.25% | 98.51% | 97.81% | 99.24% | 99.44% | 97.73% | 99.23% | 99.43% | 97.79% | 99.23% | 99.43% |
| | | F-score | 92.80% | 97.79% | 98.78% | 92.97% | 97.84% | 98.82% | 92.73% | 97.77% | 98.79% | 98.25% | 99.40% | 99.55% | 98.18% | 99.39% | 99.55% | 98.25% | 99.39% | 99.54% |
| | | True positive | 3172347 | 3205898 | 3207358 | 3172082 | 3205807 | 3207242 | 3170755 | 3206001 | 3207353 | 3181144 | 3207111 | 3207908 | 3180305 | 3206997 | 3207905 | 3181142 | 3207056 | 3207920 |
| | | False positive | 8291 | 1522 | 996 | 8325 | 1577 | 1046 | 8438 | 1595 | 1021 | 6050 | 1269 | 927 | 5969 | 1237 | 888 | 6083 | 1255 | 889 |
| | | False negative | 37909 | 4359 | 2899 | 38175 | 4450 | 3015 | 39502 | 4256 | 2904 | 29113 | 3146 | 2349 | 29951 | 3260 | 2352 | 29115 | 3201 | 2337 |
| | SNPs | Precision | 99.74% | 99.95% | 99.97% | 99.74% | 99.95% | 99.97% | 99.73% | 99.95% | 99.97% | 99.81% | 99.96% | 99.97% | 99.81% | 99.96% | 99.97% | 99.81% | 99.96% | 99.97% |
| | | Sensitivity | 98.82% | 99.86% | 99.91% | 98.81% | 99.86% | 99.91% | 98.77% | 99.87% | 99.91% | 99.09% | 99.90% | 99.93% | 99.07% | 99.90% | 99.93% | 99.09% | 99.90% | 99.93% |
| | | F-score | 99.28% | 99.91% | 99.94% | 99.27% | 99.91% | 99.94% | 99.25% | 99.91% | 99.94% | 99.45% | 99.93% | 99.95% | 99.44% | 99.93% | 99.95% | 99.45% | 99.93% | 99.95% |
| DNAscope | | | | | | | | | | | | | | | | | | | | |
| | | True positive | 443499 | 466856 | 470799 | 444329 | 467200 | 471196 | 443141 | 466836 | 471115 | 467078 | 477999 | 479118 | 466450 | 477903 | 479072 | 467027 | 478004 | 479118 |
| | | False positive | 21902 | 8738 | 6002 | 21250 | 8438 | 5540 | 21807 | 8721 | 5777 | 5615 | 1617 | 1194 | 5775 | 1648 | 1217 | 5460 | 1595 | 1195 |
| | | False negative | 37767 | 14409 | 10466 | 36938 | 14066 | 10070 | 38121 | 14429 | 10150 | 14187 | 3268 | 2149 | 14815 | 3365 | 2194 | 14238 | 3265 | 2149 |
| | InDels | Precision | 95.29% | 98.16% | 98.74% | 95.44% | 98.23% | 98.84% | 95.31% | 98.17% | 98.79% | 98.81% | 99.66% | 99.75% | 98.78% | 99.66% | 99.75% | 98.84% | 99.67% | 99.75% |
| | | Sensitivity | 92.15% | 97.01% | 97.83% | 92.32% | 97.08% | 97.91% | 92.08% | 97.00% | 97.89% | 97.05% | 99.32% | 99.55% | 96.92% | 99.30% | 99.54% | 97.04% | 99.32% | 99.55% |
| | | F-score | 93.70% | 97.58% | 98.28% | 93.85% | 97.65% | 98.37% | 93.67% | 97.58% | 98.34% | 97.92% | 99.49% | 99.65% | 97.84% | 99.48% | 99.65% | 97.93% | 99.49% | 99.65% |

**Supplementary Table 1S. Variant calling performance of PCR and PCR-free libraries with three variant callers.** Variant calls from each library and variant caller were evaluated by the Vcfeval tool in RTGtools against the NIST truth set at high confidence regions.

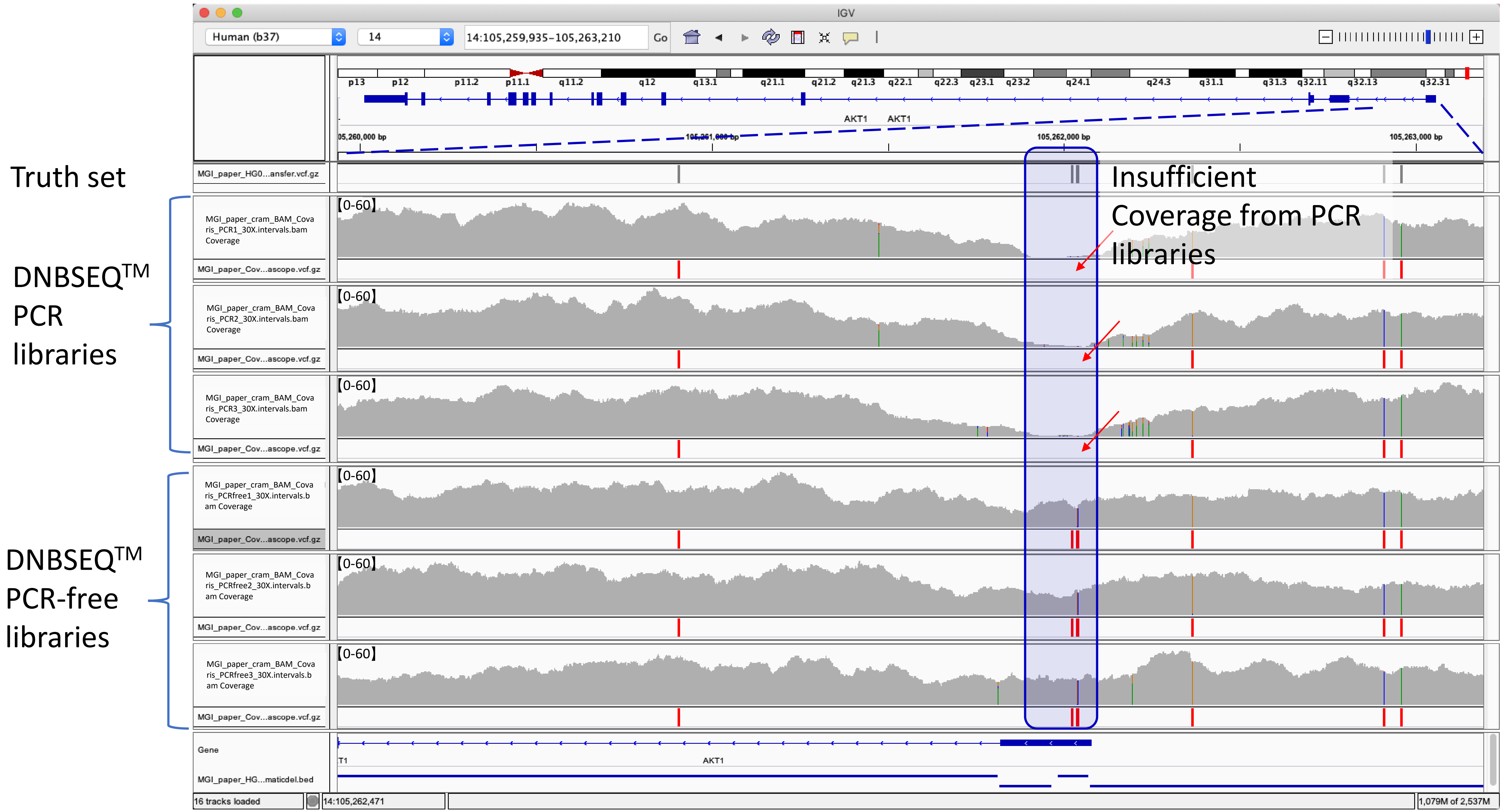


**Supplementary Figure 6S. False negative detection in DNBSEQ™ PCR libraries that may cause mis-diagnosis in clinical practice related to the ATK1 gene.** Clinical Phenotype: Cancer, Cowden syndrome 6; All PCR libraries failed to detect Chr14: 105262025 T to TC insertion at Exon1 UTR and Chr14: 105262041 CG to GC SNP at Exon1 UTR.

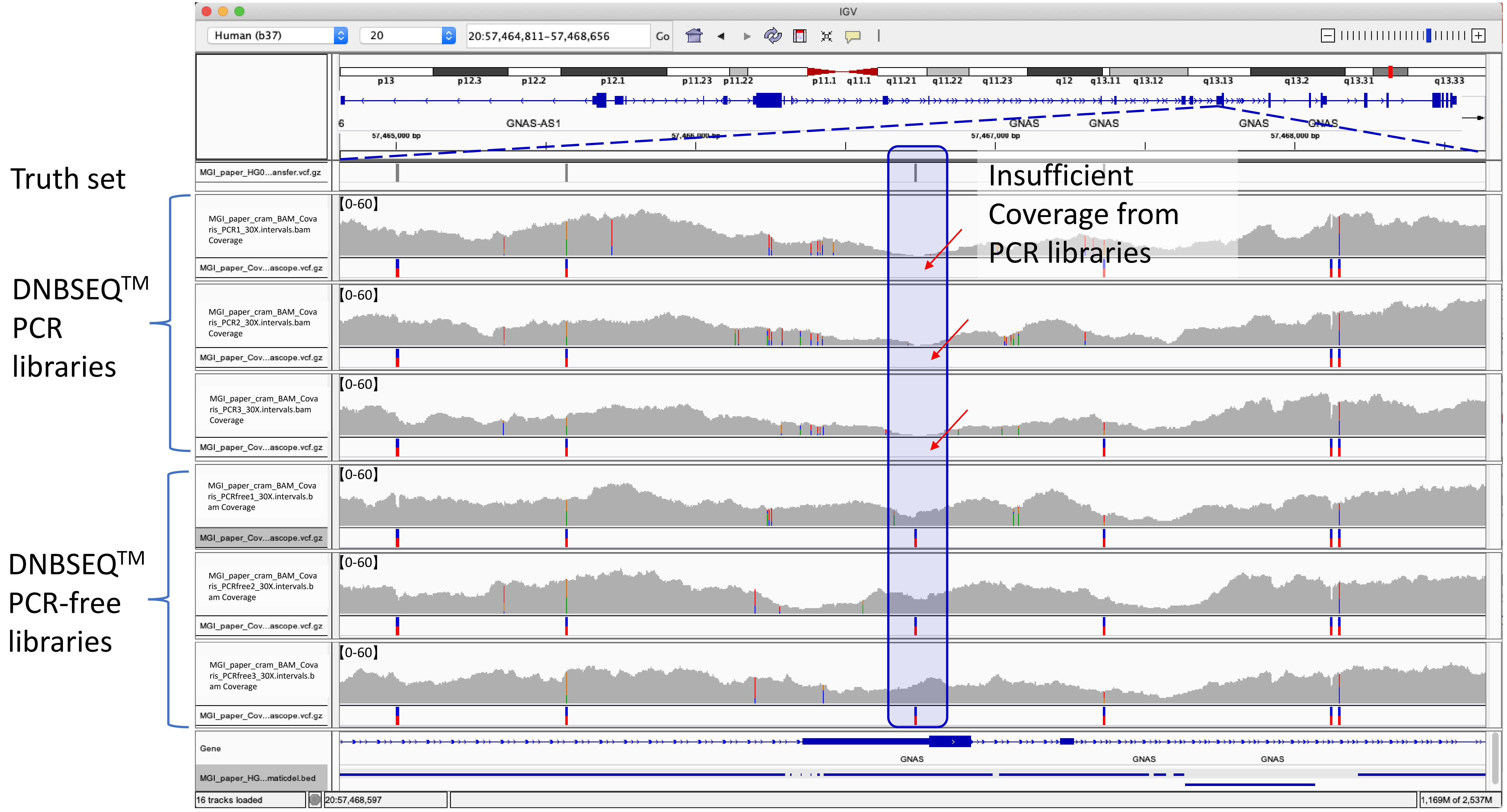


**Supplementary Figure 7S. False negative detection in DNBSEQ™ PCR libraries that may cause mis-diagnosis in clinical practice related to the GNAS gene.** Clinical Phenotype: Pseudohypoparathyroidism; All PCR libraries failed to detect Chr20: 57466734 C to CCCG insertion at Exon1 UTR.

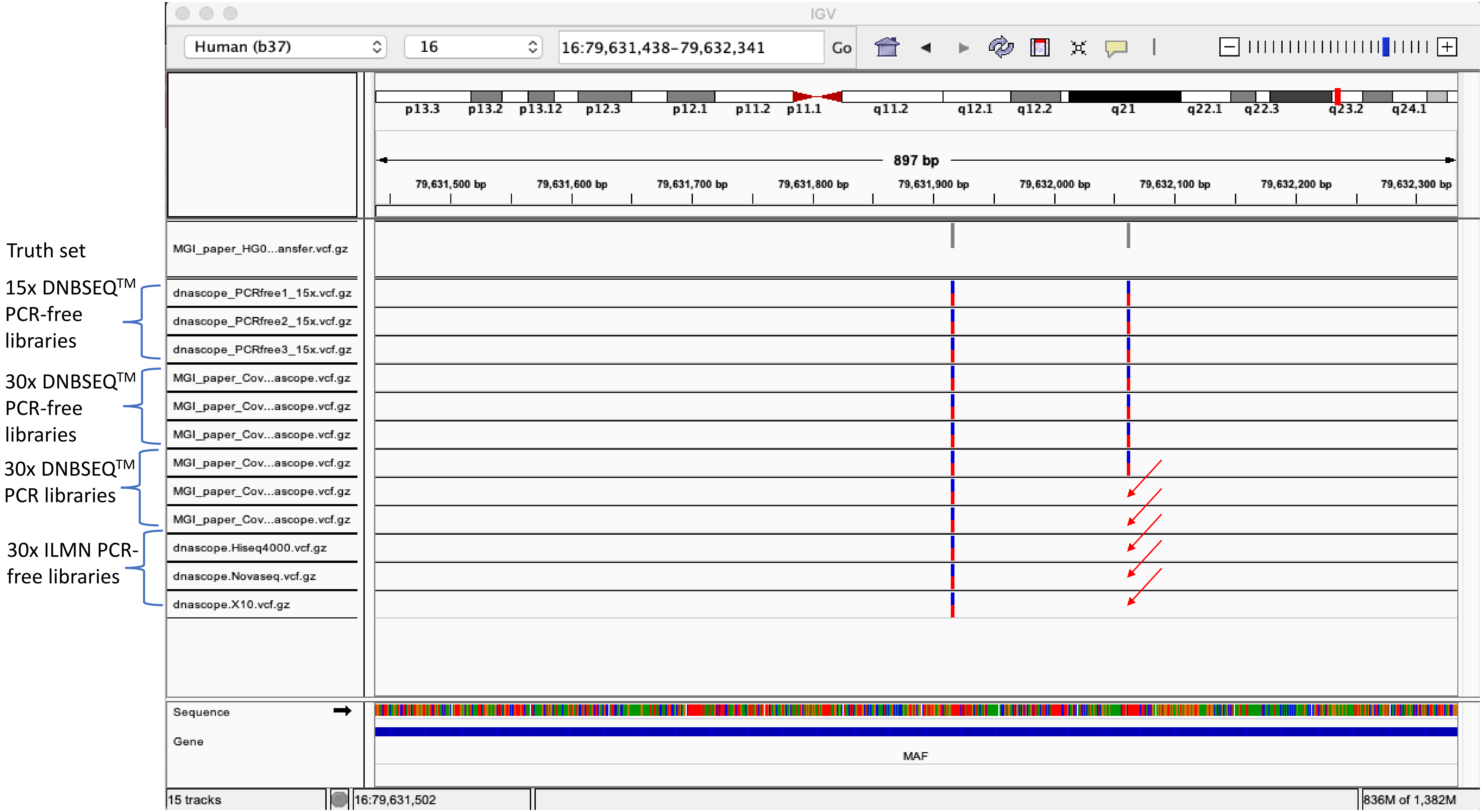


**Supplementary Figure 8S. False negative detection in 3 Illumina libraries and 2 DNBSEQ$^{TM}$ PCR libraries that may cause mis-diagnosis in clinical practice related to the MAF gene.** Clinical Phenotypes: Ayme-Gripp syndrome; Cataract 21, multiple types; All ILMN libraries failed to detect Chr16: 79632062 C to CT insertion at Exon 1, which was detected by all DNBSEQ$^{TM}$ PCR free libraries at even 15x depth.
